# Pre-sleep experiences shape neural activity and dream content in the sleeping brain

**DOI:** 10.1101/2023.07.29.551087

**Authors:** Deniz Kumral, Jessica Palmieri, Steffen Gais, Monika Schönauer

**Affiliations:** Institute of Psychology, Neuropsychology, University of Freiburg, Freiburg im Breisgau, Germany; BrainLinks-BrainTools, University of Freiburg, Freiburg im Breisgau, Germany; Institute of Medical Psychology and Behavioral Neurobiology, University of Tübingen, Tübingen, Germany; Bernstein Center Freiburg, University of Freiburg, Freiburg im Breisgau, Germany

**Author notes:** M. Schönauer, Institute of Psychology, Neuropsychology, University of Freiburg; Freiburg im Breisgau, Germany, Engelbergerstraße 41, Freiburg im Breisgau, 79106, Germany, D. Kumral, Institute of Psychology, Neuropsychology, University of Freiburg; Freiburg im Breisgau, Germany, Engelbergerstraße 41, Freiburg im Breisgau, 79106, Germany.

**Keywords:** sleep, EEG, memory reactivation, dreams, REM sleep, representational similarity analyses

## Abstract

Dreams incorporate recent experiences, and memory-related brain activity is reactivated during sleep, suggesting that dreaming, memory consolidation and reactivation are tightly linked. We devised a paradigm to investigate whether memory reprocessing during sleep contributes to dreaming. Participants listened to different audiobooks before falling asleep, introducing dissimilar experiences to be processed at night. We show that audiobook content was reprocessed at the neural level using multivariate pattern analyses. Brain activity during rapid eye movement sleep, particularly in the beta range, carried information about the audiobook. While the amount of neural reinstatement did not correlate with memory retention, global beta power during REM sleep was associated with better memory performance. Moreover, blind raters could determine which audiobook participants had studied based on dream reports. Participants who dreamt of the audiobook also showed stronger neural reinstatement. Reprocessing of pre-sleep experiences during sleep may thus shape our brain activity, our dreams, and our memories.

## Introduction

Sleep is an active state during which the brain processes new experiences in service of long-term memory storage ^1,2^. Dreams let us relive aspects of daytime experience ^3–5^, and neuronal replay during sleep strengthens and transforms recent memories ^6–10^. It has been proposed that the fragments of daytime episodes that resurface in dreams could reflect the neural reactivation of those experiences ^4,11,12^. Whether the integration of memories into dreams depends on their neural reactivation and is thus instrumental to memory consolidation, however, has not been established.

Converging evidence demonstrates spontaneous reactivation of newly encoded memories in the sleeping brain ^6^. Such experience-dependent reactivations have been observed in both animals and humans. In rodents, neural activity patterns in the hippocampus and neocortex during NREM (non-rapid eye movement) sleep ^7,13^ and REM sleep ^10,14^ reflect pre-sleep experience. Similarly, spontaneous human brain activity in both REM ^15^ and NREM sleep ^6,9,16^ reflects previously encoded information and benefits later memory performance. Furthermore, externally inducing reactivation of a previous memory task via auditory or olfactory cues during sleep boosts memory retention in REM and NREM sleep ^17–19^, suggesting a functional role of reactivation in sleep-dependent memory consolidation.

Interestingly, these findings parallel evidence showing learning-related events resurfacing in dreams ^3^. It has been shown that extensive pre-sleep activities, like playing Tetris for several hours, influence hypnagogic imagery at sleep onset ^20^. Another study observed that the content of sentences presented in an intensive study session before sleep was incorporated more often into dreams than that of other sentences ^21^. Dreaming of a learning task, such as navigating a virtual maze, can also improve participants’ performance on later memory tests ^22,23^. These findings have led to the proposal that memory reprocessing during dreams may support memory consolidation during sleep. However, the mechanisms underlying dream incorporation and whether this is driven by neural reinstatement of learning-related activity patterns remain unclear.

We devised a novel experimental paradigm to investigate whether pre-sleep experiences are processed during sleep, both in neural activity and dreaming (see Materials and Methods). Specifically, we were interested in whether the content of an audiobook listened to before sleep could be detected in both brain activity and the dream reports we collected throughout the night. Studies quantifying memory incorporation in dreaming have faced difficulty in statistically validating whether dream incorporation occurs (but see ^24^). We address this issue by systematically manipulating the content of pre-sleep learning material, and assessing its impact on the content of dream narratives. Blind raters were tasked with judging, for each dream collected during the night, which audiobook was listened to beforehand. Concordance analyses between the actual and the rated audiobook conditions then allow us to objectively quantify whether information pertaining to the audiobook is present in the dream reports. Participants were presented with one of four different audiobooks while falling asleep, aimed at introducing dissociable brain activity as well as dreams during the ensuing sleep period. After sleeping for one 90-min sleep cycle, the participants were awoken to report their dreams and retrieve the previously presented audiobook content (**Fig. 1A**). This procedure was repeated multiple times during the night, with high-density EEG recorded throughout the experiment. We can thus also assess systematic differences in neural activity induced by the experimental manipulation (see ^6^).

**Fig. 1.**
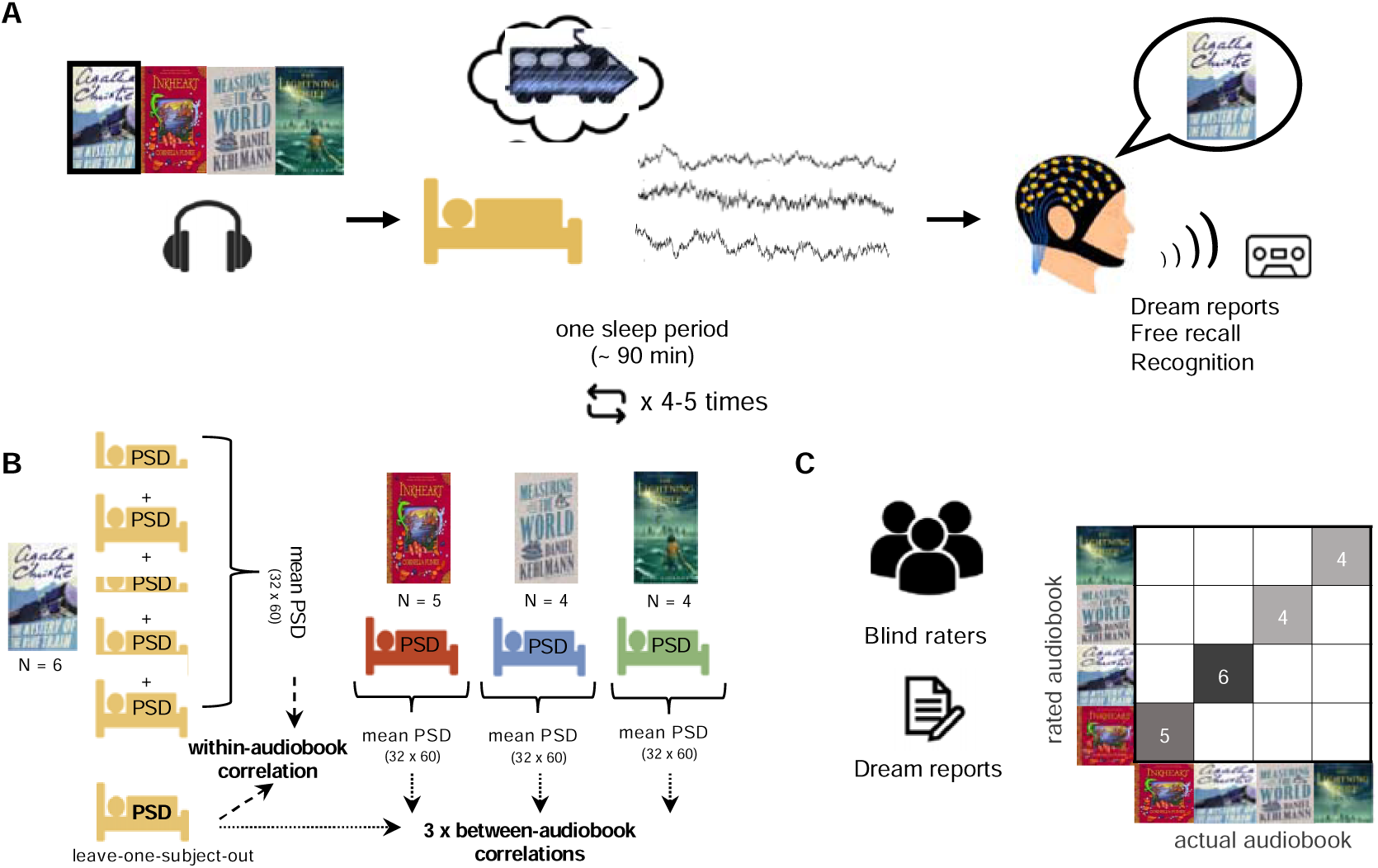
Experimental procedure. **A.** During encoding, participants were presented with one of four audiobooks (e.g., “The Mystery of the Blue Train” by Agatha Christie). While participants fell asleep, EEG was recorded using active electrode 128-channel EEG. After 90 min of sleep, participants were awoken. First, they were asked for their dreams. Next, they then had to freely recall the audiobook content they had listened to while falling asleep. Finally, they indicated which sections of the audiobook they recognized. The whole procedure was repeated up to five times over the night. **B.** We assessed representational similarity between participants using power spectral density (PSD) within and between audiobook conditions in a leave-one-subject-out approach. The within-audiobook correlation was calculated as the Spearman correlation between the average sleep PSD (dimensions: 32 channel x 60 Hz) of one individual and the average PSDs of all remaining participants in the same audiobook condition. The between-audiobook correlations were computed between the average PSD of the left-out subject and the averaged PSDs of all participants in each other audiobook condition, respectively. Results of all subjects were averaged for all possible within- and between-audiobook correlations. Finally, we computed Δcorr as the difference between within- and between-condition correlations. A higher Δcorr indicates higher similarity of neural activity between subjects in the same audiobook condition and is a measure for audiobook reinstatement in sleep EEG**. C.** We assessed whether the specific content of an audiobook is reprocessed in subsequent dreams: three blind raters were presented with isolated, anonymized dream reports and asked to judge which audiobook each participant had listened to before this dream.

We predicted that the narrative of the audiobooks would not only shape brain activity but also the content of the dreams our participants experience during sleep. We used representational similarity analysis to investigate whether spontaneous electrical brain activity during sleep holds information about the content of the recently encoded audiobook narratives ^25^ (**Fig. 1B**, see Materials and methods). Representational similarity analysis can detect content-specific neural processing in both wakefulness and sleep ^25–27^ and particularly lends itself to multivariate analyses on more than two stimulus conditions. If the previous audiobook has had an impact on brain activity during subsequent sleep, brain activity patterns during sleep periods of participants who listened to the same audiobook should be more similar than brain activity patterns during sleep periods of participants who listened to different audiobooks. To further assess whether the specific content of an audiobook is reinstated in subsequent dreams, three blind raters were asked to indicate which audiobook each participant had listened to before sleep, based on dream reports collected during the night (**Fig. 1C**). Crucially, we hypothesized that if neural reactivation shapes the content of our dreams, we should observe a stronger neural processing signal in those participants who dreamt of the audiobook. Despite a comparably small sample size (67 sleep segments from 19 participants, 50 dream reports), this study thus aims to establish proof-of-concept that naturalistic learning material presented before sleeping can influence both dream content and neural processing during sleep.

## Results

### Pre-sleep experiences shape neural activity during sleep

To test whether our experiences also shape neural activity in the sleeping brain, we investigated whether spontaneous electrical brain activity during sleep holds information about the previously played audiobook. For this, we assumed, based on previous findings, that brain activity that is influenced by similar experience will have similar features ^6,25^. We therefore extracted the spatial pattern of brain oscillatory activity in different frequency bands (power spectral density, PSD) from sleep EEG, for REM and NREM. Because PSD differs greatly between sleep and wakefulness, electrophysiological activity cannot be directly compared between these states. We thus used between-subject analyses to compare PSD from the same sleep states and correlated brain activity patterns between participants in a representational similarity analysis (RSA, **Fig. 1B**) ^6,25^. We found that during REM sleep, brain activity was more similar among participants if they had listened to the same audiobooks than if they had listened to different books. Calculating permutation statistics by performing the same analyses with the effect of the audiobook removed by shuffling the condition labels across participants confirmed that the previous experience systematically shaped brain activity during REM sleep (Δcorr = 0.063, *P* = 0.039, **Fig. 2A**). We did not find similar evidence for memory reprocessing during NREM sleep (Δcorr = -0.052, *P* = 0.785, **Fig. 2B)**. Based on previous evidence demonstrating reactivation during slow wave sleep (SWS) ^28^, we further explored whether we can detect memory reprocessing in this sleep state. Consistent with our findings in NREM sleep, we did not observe evidence supporting memory reprocessing in SWS (Δcorr = -0.04, *P* = 0.700). The failure to detect memory reprocessing in NREM sleep states may be partially due to the higher variability in PSD between participants than in REM sleep (for a similar pattern of results, see^6,25^). Finally, we repeated the RSA only in participants who reported a dream experience (N=15, 42 sleep segments). While our results indicated a similar direction for memory reprocessing in REM sleep (Δcorr = 0.049, *P* = 0.134), we found no evidence of reprocessing in NREM sleep (Δcorr = -0.019, *P* = 0.739). To further evaluate whether reprocessing occurs uniformly over time, we divided the night into early and late phases by averaging the first two and last two sleep segments for each individual. While evidence for audiobook reprocessing during REM in the second half of the night was high (Δcorr = 0.04, *P* = 0.05, averaged across N=15 individuals), we were unable to perform the same analysis in the early phase of sleep due to insufficient data, as the number of individuals experiencing REM sleep was not sufficient. However, we replicated our findings on audiobook reprocessing during late-night REM periods, which contained information pertaining to the previous learning experience. To assess the robustness of our findings, we repeated our main analyses using two variant approaches to RSA data processing, confirming the results (**see Fig. S1, S2**). Information about previous learning content can thus be detected during REM sleep, demonstrating that pre-sleep experiences are reinstated at the neural level. To assess whether brain activity during sleep retained content-specific features of pre-sleep experience, we conducted RSA on PSD data, comparing patterns from wakeful listening to sleep. Our classification analyses did not yield significant results in either REM (*P* = 0.55) or NREM sleep (*P* = 0.284), suggesting no discriminability of audiobook-related neural patterns across wake and sleep states in the present dataset.

**Fig. 2.**
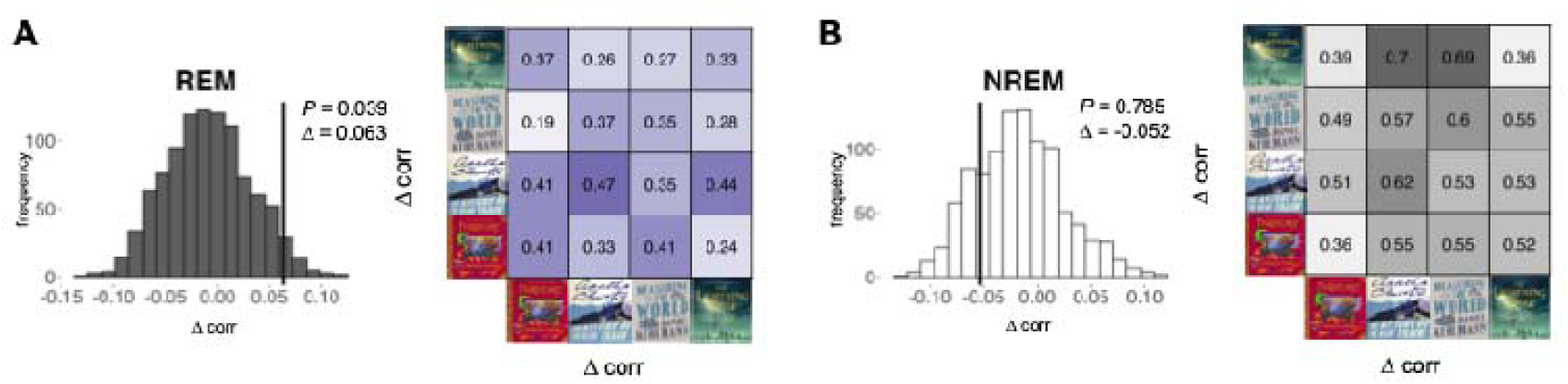
Neural reinstatement in NREM and REM sleep. It has previously been shown that brain activity patterns induced by stimulation are similar across subjects ^6,25^. Neural reinstatement is therefore measured as a higher similarity of brain activity in sleep in participants listening to the same audiobooks before sleep as compared to participants listening to different audiobooks (difference in Fisher’s z transformed correlations, Δcorr). Histograms: permutation distributions of within-between differences (Δcorr); vertical lines: observed within-between differences. **A.** Brain activity during REM sleep was informative about which audiobook participants listened to before sleep: participants who listened to the same audiobook had more similar REM activity patterns than participants who listened to different audiobooks (Δcorr = 0.063, *P <* 0.039), **B.** but not NREM sleep (Δcorr = -0.052, *P =* 0.785). The representational similarity matrices for REM and NREM show within-audiobook (diagonal) and between-audiobook (off-diagonal) correlation values (REM: within-audiobook correlation *M* = 0.398, between-audiobook correlation *M* = 0.334; NREM: within-audiobook correlation *M* = 0.491, between-audiobook correlation *M* = 0.554). Note that a total of 67 awakenings from 19 participants entered the representational similarity analyses.

### Higher frequency EEG activity reflects memory reinstatement during REM sleep

To determine which aspects of the neural signal contribute to memory reprocessing in REM sleep, we performed a frequency-of-interest (FOI) analysis on EEG activity in different frequency bands. We removed all audiobook condition information from specific EEG frequencies in the power spectrum between 0.05 and 30 Hz, by randomly re-assigning these parts of the signal between participants. If this procedure significantly lowers similarities between participants who listened to the same audiobook, the respective frequency band contains information about the narrative content and thus reflects the processing of the audiobook during sleep. Our results show that high frequency beta activity (18–30 Hz) is critically involved in the reprocessing of audiobook content during REM sleep (*P =* 0.006, **Fig. 3A**). We further confirmed this finding by performing RSA on brain activity from the FOI range only. Again, only activity in the 18–30 Hz beta band contained information about the previously presented audiobook (see **Fig. S3**). We thus suggest that high-frequency brain activity could serve as a neural fingerprint of memory reinstatement in human REM sleep.

**Fig. 3.**
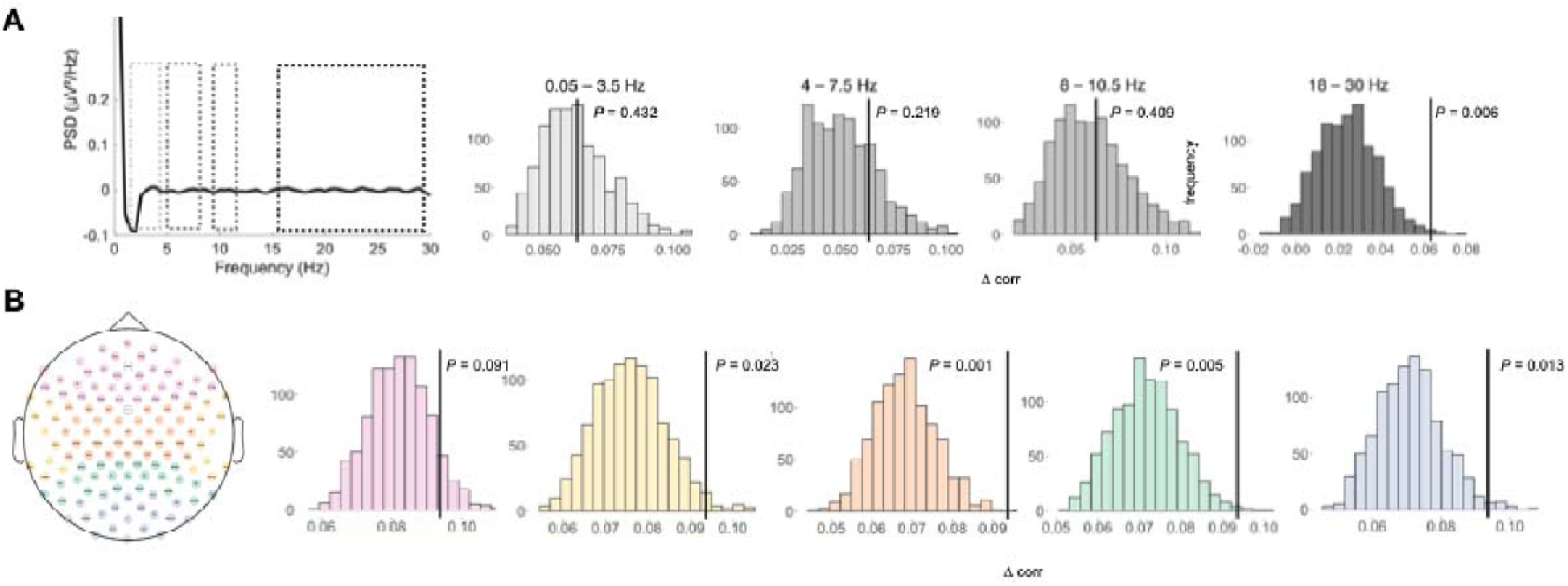
Frequency- and region-specific neural reinstatement during REM sleep. **A.** To assess the contribution of the different EEG frequency bands to memory processing in REM sleep, we shuffled the frequency ranges of interest between participants in the representational similarity analyses (RSA). This procedure thus removed the association with the audiobook condition from these parts of the data while keeping the correct condition labels intact for all other frequency ranges. If informativeness is removed from REM sleep beta activity (18–30 Hz), the audiobook condition can no longer be discerned equally well from sleep EEG (*P <* 0.006). Neural reinstatement is thus strongest in REM-sleep high-frequency beta activity. Note that permutation distributions for searchlight analyses are not centered on 0 because brain activity in other frequency ranges may also hold partial information about audiobook content. The observed Δcorr remains constant at the value reported above (Δcorr = 0.063). **B.** Region of interest analysis on frontal (pink) temporal (yellow) central (orange) parietal (green) and occipital (blue) electrodes. Electrodes over all brain regions except for frontal channels carry significant information about the content of the previously studied audiobook (p<0.05). Note that a total of 67 awakenings from 19 participants entered the RSA.

To test whether individual brain regions are particularly involved in memory processing during REM sleep in beta-frequency range, we implemented a similar searchlight approach, described above. We first grouped the EEG channels into six regions of interest (ROI) (**Fig. 3B**). By removing information about the audiobook information from six ROIs in the beta frequency range (18–30 Hz), we tested whether this lowers the similarity patterns. Our results indicate that beta activity recorded over each ROI, except the frontal cortex, was important for identifying the audiobook content (all *P* < 0.05, **Fig. 3B**), suggesting that memory reprocessing occurs across a distributed network of brain regions.

### Linking neural memory reprocessing and memory retention

To examine whether memory processing during sleep is associated with retention of the audiobook’s content, we correlated free recall and recognition performance for passages from the audiobook with the degree of memory reprocessing and the strength of oscillatory activity features that carried information about audiobook content (18–30 Hz). While we did not observe a significant correlation between the strength of reprocessing — measured as the match of an individual’s PSD with the PSD template for that audiobook condition — and free recall (r = -0.13, *p* = 0.60) or memory recognition (r = 0.08, *p* = 0.70; **Fig. 4A**), beta activity was positively correlated with free recall performance (r = 0.55, *p* = 0.01) and recognition memory (r = 0.45, *p* = 0.04; **Fig. 4B**). This indicates that while neural reactivation *per se* did not predict memory retention, elevated beta power during REM sleep correlated with better retention of the narrative content over time. This suggests that neural reactivation, which was related to beta activity or a separate physiological process modulated by beta oscillations, contributed to successful memory consolidation. Beta activity during sleep has previously been associated with the ability of participants to recall dream content^29^. Indeed, we similarly found that increased beta power during REM sleep is associated with a higher word count in dream reports (r = 0.60, *p* = 0.007). As beta activity in REM sleep may reflect a shared cognitive mechanism underlying both dream recall and the accessibility of recently encoded material, we further implemented mediation analysis to test whether general recall ability (i.e., the number of words in the dream report) mediates the association between beta power and free recall performance. Although we observed that the direct effect (c’) of beta activity on audiobook recall was no longer significant (*b*=0.35, SE=0.25, t(16)=1.36, *p*=0.18), the mediation analysis was not significant, as the bootstrapped indirect effect confidence interval included zero (*b*=0.23, SE = 0.3, with a 95% CI of [−0.07, 0.85]).

**Fig. 4.**
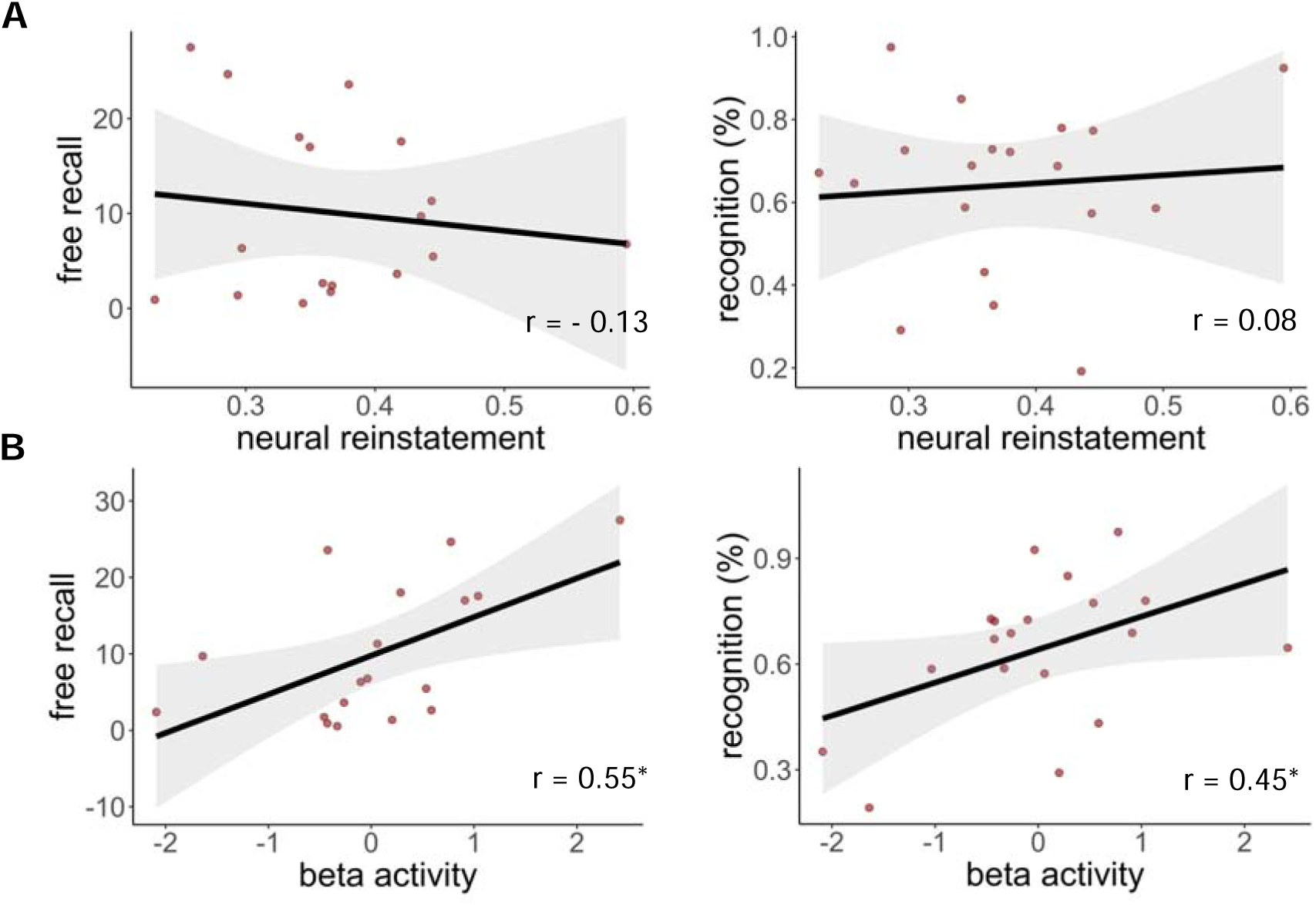
Linking neural activity and memory. Using Pearson’s correlations, we related memory performance with neural reinstatement and averaged beta activity (18–30 Hz) in REM sleep with later memory performance. **A.** We did not observe relation of reprocessing strength in REM sleep with free recall (r = -0.13, *p* = 0.6, n=18) or memory recognition (r = 0.08, *p* = 0.70, n=19), **B**. Higher activity in the beta frequency range in REM sleep correlated with both higher free recall (r = 0.55, *p* = 0.01, n=18) and better recognition memory (r = 0.45, *p* = 0.04, n=19). *p < 0.05.

### Pre-sleep experiences shape dream content during sleep

We further predicted that the narrative of an audiobook should shape the content of ensuing dreams. To assess whether the specific narrative of an audiobook is reactivated in dreams, three independent and blind human raters were asked to judge which audiobook each participant listened to before having a particular dream **(Fig. 1C)**. The blind raters were able to determine which audiobook the participants had listened to before sleep with above-chance-level accuracy, based solely on their dream reports. We found significant concordance between the rated book and the actual audiobook that participants listened to (analysis using individual ratings: ĸ = 0.107, z = 2.29, *p* = 0.02, percent correct ratings = 32.9%, **Fig. 5A**; confirmation when combining the ratings across raters for n=39 dreams: ĸ = 0.174, z = 1.78, *p* = 0.07, percent correct ratings = 38.2%). Please note that these ĸ values describe the agreement of ratings given by three independent raters, i.e., the rated audiobook condition, with the actual audiobook condition of each dream (ground truth) in a four-way forced-choice decision. Thus, these values contain various sources of noise, including participants not having dreamt of anything relating to the audiobook, not reporting potential aspects of the dream related to the audiobook, and raters not recognizing aspects pertaining to the audiobook (e.g., because of imperfect memory for the source material or attentional lapses), such that the ĸ values are expected to be low in magnitude. For similar reasons, the agreement among the three raters was low yet significant (16.3%, ĸ = 0.172, *p* = 0.0003), indicating that they relied on the same sources of information in the dream reports when providing their ratings. We further tested whether dreams collected after awakenings from REM and NREM sleep equally carried experience-driven information. Notably, it was REM dreams in particular that contained information about the previous audiobook. Raters were able to determine the correct audiobook based on dream reports collected after REM sleep awakenings (analysis on individual ratings: ĸ = 0.343, z = 2.88, *p* = 0.003, percent correct ratings = 54.2%, **see Fig. S4A**). However, they were unable to do so above chance level based on dream reports collected after NREM sleep awakenings (analysis on individual ratings: ĸ = 0.08, z = 1.43, *p* = 0.154, percent correct ratings = 31.5%, **Fig. S4B**). Note that REM sleep reports were predominantly collected later during the night, when participants have had longer exposure to the audiobooks. Time of night and amount of audiobook exposure could both be important drivers of dream incorporation, other than sleep stage. In summary, we therefore conclude that pre-sleep experiences impact our dreams during later sleep.

**Fig. 5.**
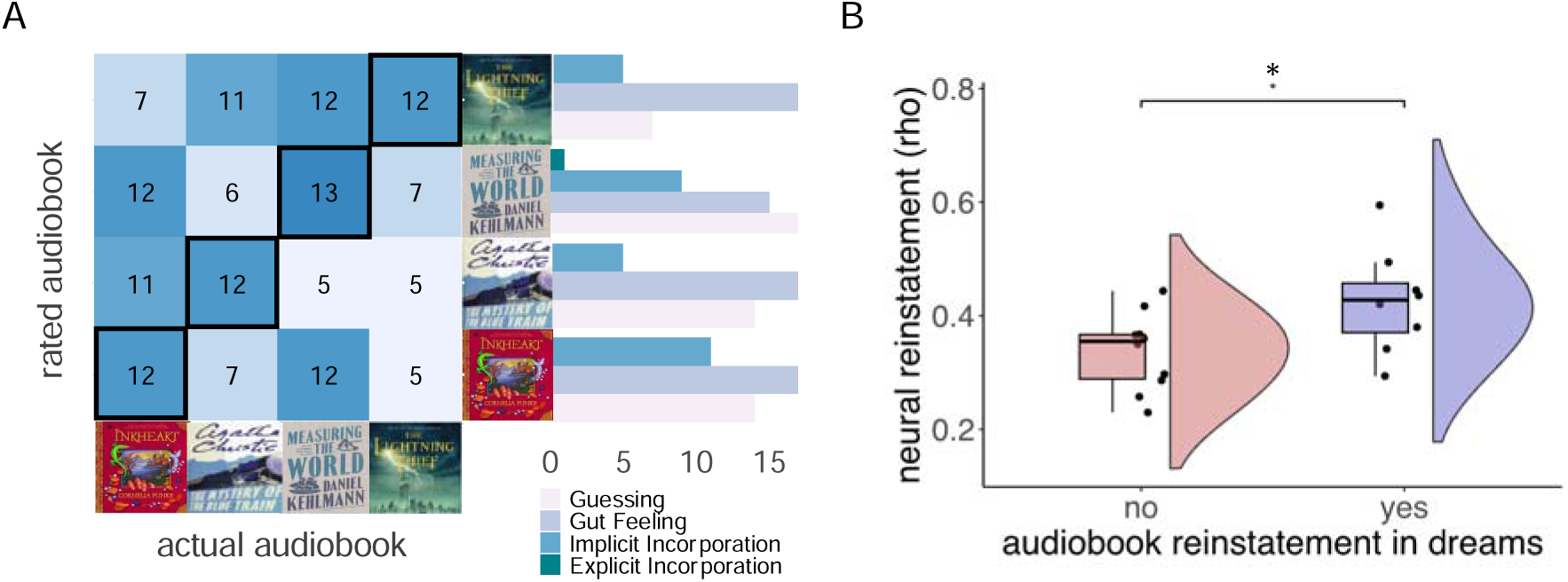
Classified dream content and neural reprocessing. **A.** Blind raters were able to judge with better than chance accuracy which audiobook the participants had listened to before sleep based on transcripts of the dream reports (ĸ = 0.107, z = 2.29, *p* = 0.02, percent correct ratings = 32.9%, based on 149 dream ratings by three raters). Since many dreams did not include direct audiobook incorporations, the raters most frequently based their judgements on *gut feeling*. Note that also many *implicit incorporations* occurred, whereas *explicit incorporations* are rare. **B.** A two-sample t-test revealed that the amount of neural reactivation during REM sleep was higher in participants who had incorporated audiobook information into their dreams (*t*_16_ = 2.4, *p* = 0.03, Cohen’s d = 1.10). \**p* < 0.05.

### Neural memory reprocessing shapes dream content

If neural reactivation shapes the content of our dreams, we should observe a stronger neural reprocessing signal in participants who dreamt of the audiobook. To test this hypothesis, we compared the neural reprocessing strength between participants whose dreams contained information about the audiobook condition and participants whose dreams did not reflect the audiobook. In addition to assigning the likeliest audiobook to a dream report, dream raters quantified the amount of information in participants’ dreams pertaining to the audiobook (see Methods). Based on this score, we identified dream incorporators and dream non-incorporators. As predicted, we observe a higher reprocessing strength in participants who dreamt of the audiobook (*t*_16_ = 2.4, *p* = 0.03, Cohen’s *d* = 1.10, **Fig. 5B**). The content of our dreams is thus at least partially linked to experience-related neural activity, demonstrating that pre-sleep experiences effectively shape both brain and cognitive activity during sleep.

## Discussion

By manipulating the content of pre-sleep learning material, we could determine which audiobook someone had studied before going to bed, based on their dream reports. Naturalistic stimulus material like encoding the narrative of an audiobook may be particularly effective for inducing dream incorporation. Stories elicit deep, semantic processing, and often affect us emotionally, resulting in a sustained influence on spontaneous thought ^30^. Consistently, pattern analyses of spontaneous electrical brain activity revealed neural reprocessing of audiobook content during REM sleep. Activity in the high-frequency beta range in REM sleep was further associated with memory retention. When participants dreamt of the audiobook during the night, we also found stronger reinstatement at the neural level. That our waking experiences play an important role in shaping our dreams is in line with suggestions that dreams may reflect memory reactivation and consolidation during sleep ^4,22,31^. To date, only a small number of empirical studies have linked memory processing with dreaming (for a detailed review, see ^3,28^). Verbal material presented prior to sleep, for example, has been found to be incorporated into dreams during both NREM and REM sleep, and more frequently than comparable material that was not presented beforehand ^21,24^. Our study provides a first indication that memory reprocessing at the neural level results in the reappearance of such learned content in subsequent dreams, supporting the idea that dreams may partially reflect memory consolidation during sleep.

### Memory reprocessing of complex narratives in REM sleep

Brain activity patterns during REM sleep held information about which audiobook our participants had listened to before falling asleep. The majority studies investigating memory reactivation during sleep have focused on the role of NREM sleep. Our results indicate that pre-sleep experiences like listening to a complex narrative are similarly reprocessed during REM sleep. Our results agree with early behavioral studies suggesting REM sleep actively facilitates memory consolidation ^32,33^. This idea is further supported by reports of increased REM sleep density after periods of learning ^34,35^, a positive association between the amount of post-learning REM and performance outcomes ^36,37^, as well as more recent findings that REM sleep not only reinforces memories ^38^ but may also promote hippocampal reorganization by enhancing the differentiation of memory representations ^39^.

In line with these findings, previous studies provide noteworthy evidence of spontaneous reactivation of learning experiences also during REM sleep^6,10,15,35,40–42^. In the rat hippocampus, neuronal firing patterns observed during path running are replayed in subsequent REM sleep ^10^. Recent research suggests a role of REM sleep in the consolidation of novel experiences ^35^ through both neural assembly reactivation and sequential replay of learning-related neuronal activity ^41^. Consistent with this idea, the distribution of cerebral activity during REM sleep is modified by previous learning experience in humans ^15^.

Whether memories are processed during NREM or REM sleep may depend on the type of learning task. Particularly complex learning material may benefit from processing in REM sleep ^33,43^. Moreover, recall of learning material with similar complexity to ours (short stories) was impaired by REM sleep deprivation ^33^. However, the function of REM sleep for memory is still far from being understood (e.g., for studies linking REM sleep and forgetting see ^44,45^). Our results suggest that REM sleep is involved in both the reprocessing and the consolidation of complex narratives.

While we observed neural reprocessing in REM sleep, we did not detect significant reprocessing of memories during NREM sleep. Oscillatory activity in NREM sleep is more variable than in REM sleep (e.g., spindles, slow oscillations, mixed theta rhythms in NREM sleep), and firing behavior of neurons differs greatly between these sleep stages ^10^. This variability could potentially have prohibited equally efficient detection of learning-related activity in NREM compared with in REM sleep. Indeed, a previous study using a similar approach as we applied here was also better at decoding memory content from REM sleep than from NREM sleep ^6^. We therefore interpret our null findings in NREM as a possible consequence of higher variability in the signal rather than an absence of reactivation. Moreover, REM sleep occurs predominantly towards the end of the night, when participants have had longer exposure to the audiobook. This may have increased the audiobook-related signal in neural activity and thus facilitated audiobook discrimination in REM compared with NREM sleep.

### Detecting reprocessing of complex narratives during sleep in humans

Demonstrating wake-to-sleep reactivation in humans directly using scalp EEG is challenging due to differences in brain activity between sleep and wake states, including changes in PSD. Previous studies have shown a recurrence of learning-related activity patterns observed in the waking brain after the presentation of TMR cues ^46,47^ and occurring spontaneously, time-locked to sleep slow oscillations ^9^ during NREM sleep. The latter work ^9^ used controlled stimuli (e.g., images) and localizer data to isolate brain patterns for wake-to-sleep reactivation analysis. In contrast, we used naturalistic and complex narratives (i.e., continuous data), making classification of EEG patterns from wake-to-sleep difficult. To overcome this, we employed a between-subject analysis approach ^6,25^, which enabled us to detect learning-related information using data recorded in the same states of consciousness, avoiding noise induced by oscillatory differences between wakefulness and sleep states. Since we do not expect reactivation to occur “verbatim”, i.e. reinstating the exact sensory signals in response to auditory inputs while listening to the audiobook, this more indirect approach also provides a higher likelihood of capturing patterns related to schematic and abstract forms of reactivation, focusing on the core events and topics of the narratives. Other studies have followed an approach similar to the one applied here, detecting distinct patterns of content processing during sleep, induced by presenting TMR cues associated with different stimulus categories in NREM sleep ^48,49^, or by manipulating the content of pre-sleep learning material ^6^.

We observe REM-dependent neural reprocessing across most scalp regions, indicating that a wide network of brain regions is active during audiobook-specific reprocessing during the night. This may include activation of sensory-processing areas ^28^, the emotion processing network, as well as the reactivation of brain regions storing memories relating to the audiobook content. Audiobooks constitute information-rich stimulus material, such that it is impossible to dissociate these information sources in the neural reprocessing signal we detected here. Future studies are needed to test which aspects of memory for naturalistic narratives are reprocessed across different brain regions during sleep.

### REM sleep beta activity, memory reprocessing, and memory consolidation

Particularly higher frequency oscillatory activity in the beta frequency range carried information about the previously presented material. Moreover, the amount of activation in the beta frequency band during REM sleep was associated with post-sleep memory recall and recognition performance. These results align with previous studies investigating the role of beta activity in cognition, including memory encoding, memory search, and also dream recall ^29,50–53^. Beta activity reaches its maximum across the night during REM sleep^54^ and increases when participants are able to report the content of their dreams ^29^. While our results assign an active role of REM sleep beta activity in memory processing, the amount of beta-related memory reprocessing was not directly related to memory retention. Instead, overall power in the beta band correlated with participants’ ability to retrieve information about the audiobook as well as the content of their dream experiences. Beta-band activity during REM sleep may reflect a state of general cortical arousal or hypervigilance ^55^ Consistent with this idea, elevated REM beta power is often observed in conditions such as insomnia ^56^ and heightened arousal ^55,57^. A difference in emotional arousal induced by the individual audiobooks may thus have allowed differentiation of the previously learned content based on brain activity during sleep.

Although the positive relation with memory retention suggests a functional role of REM beta activity in memory consolidation, REM beta activity similarly correlated with dream recall ability. The observed link between higher beta power and post-sleep memory performance may thus reflect a shared cognitive mechanism underlying both dream recall and the accessibility of recently encoded material, rather than being mediated by memory reactivation of the story content. To explore this, we tested whether dream recall— approximated by the number of words in dream reports—mediated the link between REM beta activity and audiobook recall. Although the mediation was not significant, controlling for dream recall weakened the relationship between beta activity and memory performance. REM beta activity may thus at least partially reflect a general state of heightened accessibility that supports both dream recall and memory performance, rather than selectively indexing the reprocessing of specific episodic content. Alternatively, neural reactivation may drive beta activity, memory consolidation, as well as dream generation, assigning a functional role to REM beta in memory consolidation, suggesting it could serve as a neural fingerprint of memory reprocessing in humans ^29^. Future studies should investigate the role of REM sleep beta in neural reactivation, dream generation and accessibility of mnemonic content, separately, to disentangle the independent contributions of these factors.

Finally, memory performance may have been affected by factors other than REM beta activity, such as the amount of exposure to the audiobook, the amount of sleep after audiobook encoding, and the extent of sleep disruption by the serial awakening paradigm, since these were not controlled for in our study. Despite these sources of variance, we find that REM sleep beta activity correlated with measures of memory performance and carried information about audiobook content. Future studies should control the amount of information encoded before sleep and sleep duration to support this finding.

### Memory reprocessing and dream incorporation

Blind raters were able to decide, based on the content of dream reports alone, which audiobook participants had listened to before falling asleep. This indicates that information relating to the audiobook is incorporated into dream experiences. Our findings further suggest that neural reprocessing is related to incorporating the same or related information into dreams. Participants who incorporated information about the audiobook during the night also showed stronger neural reprocessing. However, dreaming experiences also comprise many aspects that cannot be directly linked to previous waking events. Thus, disentangling the different information sources of dream content is an exciting avenue for future research. Moreover, dreaming does not always occur as a narrative, but can also manifest as simple thoughts or perceptual imagery. It has been suggested that neural replay in the sensory cortex can surface as perceptual imagery during sleep and dream states ^13^. Whether the characteristics of memory reprocessing at the neural level directly relate to how memory sources surface in dreaming, and whether the format of incorporation bears implications for their functional relevance in memory consolidation presents another exciting opportunity for further investigation.

Listening to the audiobook while falling asleep, especially the last snippets before dozing off, might not allow participants to discriminate potential dream-like experiences induced by the audiobook during sleep-onset from mentation experiences during consolidated sleep, which could explain why we found memory sources in dreaming. However, in studies assessing hypnagogic imagery, participants are usually woken up at very short time intervals during sleep onset to increase the chance of detecting the ongoing phenomenon ^20,58,59^ because memory of imagery content at sleep onset that is not retrieved upon awakening fades rapidly and can hardly be retrieved at later time points. In our study, participants were awoken after 90 minutes of consolidated sleep. This minimizes the likelihood of having reports reflecting hypnagogic experiences at times when the audiobook was still playing. Moreover, neural reprocessing of the audiobook during the last quarter of an hour of the sleep period was related to audiobook incorporation into dreams, strongly supporting the view that the reported dreams also occurred within that timeframe.

## Limitations of the study

A major limitation of this study is the comparatively small sample size, with 67 sleep segments obtained from 19 participants who listened to four different audiobooks. This limited sample size reduces the study’s statistical power, making it difficult to detect anything other than very large effects. Given this constraint, our findings should be interpreted with caution and regarded as proof-of-concept. Future studies, ideally conducted with larger sample sizes and as preregistered reports, are needed to rigorously investigate memory reprocessing in REM sleep using paradigms with naturalistic stimuli and to validate the effects observed here. Another limitation of the present study is that participants were likely aware of the study’s general aims due to the repeated dream reporting, and retrieval assessments. As full blinding was not feasible, the serial awakening paradigm may have introduced bias, potentially inflating incorporation rates or increasing the recognizability of audiobook-related dream content.

Not all dream reports will include an incorporation of the stimulus material. Forced-choice rating paradigms will lead to a higher number of guesses compared to other methods. While this allows a hypothesis test of whether information about the previous audiobook is present in dream reports, estimates of incorporation rates and which memory sources underlie these incorporations depend heavily on the blind raters’ rating style and criterion, and must thus be interpreted with caution. As dreams are likely constructed from mixed memory sources, the use of complex stimulus material like in our study could represent an additional challenge for external raters. In many instances, raters accurately classified dream reports with low confidence. It may be difficult for blind raters to detect more abstract audiobook-related references with high confidence, compared to participants themselves, leading to conservative confidence rating even in the presence of stimulus-related incorporation.

Since we used a serial-awakening paradigm^60^ combined with repeated memory encoding throughout the night, letting participants repeatedly fall asleep while listening to the audiobook, the audiobook was running for a limited time while participants were asleep. This stimulation itself may have influenced dream content independent of memory reprocessing. While lucid dreamers have been reported to incorporate highly complex semantic information into dream narratives while the information is presented ^61^, we are not aware of similar reports demonstrating that complex information presented during “regular” sleep gets incorporated into dreaming, and is remembered upon waking up after a delay. There is no data on how long after presentation external stimuli presented during sleep can influence dream experiences. Future studies could control for this confound by presenting complex information during sleep only and testing whether this leads to a similar amount of dream incorporations.

Lastly, the selected narratives that may have varied in emotional engagement, which in turn could have impacted dream incorporation and brain activity during sleep. While we did not measure subjective emotional involvement while or after listening to the story, after reporting their dreams, participants were asked to rate negative and positive affect of their conscious experiences. We found no effect of audiobook on overall dream emotionality (Palmieri et al., 2025, *under review*).

In conclusion our results demonstrate that pre-sleep experiences concurrently influence our brain activity during sleep and the content of our dreams. Moreover, if participants dreamt of the learning material, they also showed stronger neural reprocessing. This suggests a link between memory reprocessing during sleep and dream content. Our proof-of-concept study raises further questions about the memory sources of dreaming: in which form are life events recapitulated and interpreted by the sleeping brain? Does dreaming itself have a functional role in memory processing? Our results suggest an intricate interplay among our waking experiences, memory reactivation, dream content, and memory storage. Unraveling the neurophysiology of dreaming and its interaction with memory functions can give insights into how memories are reprocessed during sleep and will shed light on the mechanisms that govern the emergence of conscious experience in the sleeping brain.

## Author Contributions

Conceptualization: SG, MS Methodology: DK, MS

Investigation: MS Visualization: DK

Funding acquisition: SG, MS

Formal Analysis: DK, JP, MS

Project administration: MS, SG

Supervision: MS

Writing – original draft: DK, MS

Writing – review & editing: DK, JP, SG, MS

## Acknowledgments

We would like to express our gratitude to Susanne Kirchner-Adelhardt and Maximilian Schneider who helped with data collection. Thank you also to research assistants Eduard Stroukov and Sammy-Jo Wymer who helped with data checks.

This project was supported by Deutsche Forschungsgemeinschaft (DFG, German Research Foundation) – grant: SCHO1820/2-1, Project number: 426865207, and grant: GA730/3-1.

## STAR Methods

### RESOURCE AVAILABILITY

#### Data and code availability

- **Data**: The sleep EEG raw data —recorded after the audiobook was turned off until approximately 30-60 seconds before awakening —and all dream reports, are publicly available in DREAM: A Dream EEG and Mentation database62 (DOI: 10.60493/31mg4-mfq53).
- **Code**: All original code has been deposited at the Open Science Framework repository (DOI:10.17605/OSF.IO/2BKUS) and is publicly available. DOIs are listed in the key resources table.
- **Additional information**: Any additional information required to reanalyze the data reported in this paper is available from the lead contact upon request.

### EXPERIMENTAL MODEL AND STUDY PARTICIPANT DETAILS

#### Study Participant

20 participants (10 male) aged 20 – 30 years (25.5 ± 2.7) completed the study. They were healthy, nonsmokers, and did not ingest any alcohol, caffeine or medication other than oral contraceptives on the days of the experiment. The participants reported sleeping between 6 and 10 hours per night, had a regular circadian rhythm, and were neither extreme morning nor evening chronotypes, as measured by the Munich Chronotype Questionnaire. They had no shift work or long-distance flights during the six weeks preceding the experiment and did not have any sleep-related pathology. All participants were right-handed, confirmed by the Edinburgh Handedness Questionnaire. We only recruited frequent dreamers who reported to remember the content of one dream at least three nights per week. While there is no reason to assume sex or gender differences on our outcome measures, our study design precludes assessing the effect of gender on memory reactivation or dream incorporation during sleep.

#### Ethics Statement

The experiment was approved by the local ethics committee (Department of Psychology, Ludwig-Maximilians-Universität München, Germany, reference GA730/3-1). All participants provided written informed consent prior to the experiment. Our proof-of-concept study focused on exploring how pre-sleep experiences like listening audiobooks influence brain oscillations and dream content during sleep. Our study did not evaluate health-related outcomes.

#### Stimulus Material and Experimental Design

To investigate dreaming experience and memory retrieval, we used a serial awakening paradigm ^29^. All participants visited the sleep laboratory twice, once for an adaptation night to become familiar with the experimental procedure and environment (i.e., wearing an EEG cap), and again for the night of the main experiment. On the experimental night, participants fell asleep while listening to one of four randomly assigned audiobooks: Inkheart by Cornelia Funke, The Mystery of the Blue Train by Agatha Christie, Measuring the World by Daniel Kehlmann, or Percy Jackson & the Olympians: The Lightning Thief by Rick Riordan. Participants had not read the book or listened to the audiobook prior to the experiment. The stories were chosen from different genres and to be of different content such that the addressed topics, emotions, and narrative styles differed, with the goal of increasing dissociable information in neural activity and dream experiences. We presented the German versions of the commercially available audiobooks to participants, read by different male readers.

Participants were instructed to pay close attention to the audiobook so they could recall the content when being awoken but to fall asleep when they felt the need. Simultaneously with the audiobook a tone was presented every 15 seconds. Participants were required to press any key of a computer mouse in response to that tone to ensure they were paying attention. The audiobook was turned off once they reached consolidated stage 2 sleep and no longer responded to the tone presentations. The audio was presented via external speakers at a noise level of 45 dB ± 2.92 dB (mean ± SD), measured with a sound meter at the approximate location of the participants head in bed, ensuring that they were able to hear the presented story. Approximately every 90 minutes during the night, participants were woken by the experimenter entering the room and addressing them by name. If no wakening response occurred, they were lightly touched on the arm and addressed verbally at the same time. Participants were prewarned of this procedure. They then were asked about any conscious experiences they had during sleep and about the content of the audiobook passage they had listened to (see section Dreaming and Cognitive Measures). After these tests, they continued listening to the same audiobook from the moment it had been turned off before, while falling asleep again. The audiobook was played from beginning to end. The duration of the segment listened to before participants fell asleep ranged from 6 to 121 minutes (*M* = 29.28, *SD* = 20.44 minutes). During each consecutive awakening, the audiobook was resumed at the last section the participants listened to before they had fallen asleep previously.

Each participant was awoken up to five times. One participant had to be excluded because due to a technical problem we had no information about audiobook timing. Two participants finished their audiobook before the experimental night was over. They later continued listening to a different audiobook. Data from the two affected awakenings was excluded. All other data from a total of 67 awakenings from 19 participants entered the analysis. Further details about the experimental procedure can be found in **Fig. 1**.

##### Dreaming and Cognitive Measures

Participants were awoken and reported their cognitive experience (i.e., dreaming) in a standardized dream recall procedure, after consecutive 90-min sleep segments across a full night of sleep. After the experimenter entered the sleep chamber and woke the participant, they were asked what was going through their mind immediately before waking up, and to report any conscious experience they might have had. This question was repeated up to three times. If the participants were able to remember any dream, we instructed them that they should proceed to give a detailed report on who participated in the dream, where the dream was set, and what happened in the dream. We recorded their full dream report, asking up to three times “Can you recall more?” We then proceeded to inquire further details regarding the dream content with a custom questionnaire: Who had been part of the dream? Where had the dream taken place? What happened in the dream? What was the perspective of the dream? Did the participant experience any emotions while dreaming? What would they name as the central element of their dream? After reporting their dreams, participants also had to retrieve the audiobook content. They were first asked to freely recall what they could still remember from the previous audiobook passage they had listened to before falling asleep. These reports were recorded on a voice recorder and transcribed. A person blind to experimental conditions and the dreams reported by participants was asked to redact the free recall by removing all comments related to the experimental procedure (e.g., “this time I remember a lot,” “I cannot remember”, “I do not know”), comments about the recalled content (e.g., “what a strange name,”), comments addressed at the experimenter (e.g., “you know what I mean”), and by removing unnecessary filler words like “ummm” and repetitions. The remaining text was then redacted to short full statements describing the content of the previous audiobook section, using simple active language. These procedures were implemented to make the length of the reports respective to informational detail more comparable across participants. Word count on the redacted free recall reports divided by the time (in minutes) they had listened to the audiobook while awake was used as a measure for memory performance. Please note that we implemented no additional steps to correct for the correctness of the recalled content.

Since participants volunteered information, only correct information was recalled, with varying amount and level of detail. To test their audiobook recognition memory, participants were then aurally presented with parts of the audiobook passage they had listened to previously and asked to indicate whether they still remembered having listened to it earlier (certainly yes, do not know, certainly not). Recognition memory was calculated as the audiobook runtime of yes responses divided by the total time listened while participants were awake. All participants had listened to at least 5 minutes of audiobook before falling asleep (range from 6 to 121 minutes; *M* = 29.28, *SD* = 20.44 minutes). Please note that we did not collect recognition performance on control statements that were not presented to the participants before sleep. We thus cannot give a measure of memory specificity, but simply a hit rate.

#### EEG Recording

Sleep EEG was recorded using an active 128 channel Ag/AgCl-electrode system (BrainAmp MR with ActiCap, Brain Products, Gilching, Germany) with a 1 kHz sampling frequency and a high-pass filter of 0.1 Hz. Electrodes were positioned according to the extended international 10–20 electrode system. For sleep scoring, recordings were split into 30-s epochs and sleep stages were determined on electrodes C3/C4 according to standard rules by two independent raters. Discrepant ratings were decided by a third rater. Average sleep durations are reported in **Table S1.**

The sleep EEG raw data —recorded after the audiobook was turned off until approximately 30-60 seconds before awakening —and all dream reports, are publicly available in DREAM: A Dream EEG and Mentation database ^62^ (DOI: 10.60493/31mg4-mfq53).

#### EEG data preprocessing

We used the same preprocessing approach that has been employed to detect memory reprocessing during sleep in one of our previous publications ^6^. First, EEG data were split into 4-s trials. Artefact rejection was done in a semiautomatic process using custom MATLAB 2021a (MathWorks) scripts (channels with bad recording quality, muscle artifacts, eye movements, jumps) on these trials. Also REM periods containing eye movements artefacts were removed from the data. Since the recapitulation of eye movements has been associated with successful memory recall ^63^, and eye movements during REM show similar characteristics to those in wakefulness ^64^, eye movements during REM sleep may recapitulate patterns observed during learning, and even relate to dream content ^65,66^. We refrained from correcting eye movements by means of independent component analysis to ensure no residual activity resides in the EEG trace that might allow differentiation of the previous learning material based on eye movements alone, instead opting for a complete rejection of phasic REM. Rejected channels were interpolated using EEGLAB^67^. To compute the spectral power, we performed Fourier transformation using the Welch method by averaging over 10 Hamming windows of 2-s length with 95% overlap, resulting in smooth power spectra with a final data resolution of 0.5 Hz. To test whether memories of complex narratives are reactivated in sleep and whether this reactivation is related to participants’ dreaming experience, we analyzed neural similarity of brain activity patterns during the time before we woke participants from their sleep. The last 250 trials of each sleep period before awakening entered these analyses, corresponding to the last quarter hour of sleep (16.7 minutes) during which the reported dreams had most likely occurred. We thus also ensured that a comparable number of epochs per subject entered the representational similarity analysis (RSA). All included trials were artifact-free (Schönauer et al., 2017). The duration (in minutes) of EEG data for each participant and sleep stage is reported in **Fig. S5** and descriptive statistics of NREM and REM duration used in representation similarity analysis is shown in **Table S2.**

There is considerable variability in the literature regarding the time window used to link neural activity to dream recall. While Siclari et al. ^29^ focused on the last 20 seconds before awakening, in another study the same authors ^69^ focused on a 60-second window. We opted for a longer segment to allow more stable PSD estimates in NREM and REM sleep. Our primary goal was not to relate immediate pre-awakening features in the EEG to dream recall but to assess the discriminability of audiobook conditions in NREM and REM sleep and test for a putative link to the content of dream experiences recalled upon awakening. While this precludes specific analyses of how immediate pre-awakening features in brain activity of different sleep stages relate to dream content, it increases signal-to-noise ratio for the RSA that detects neural reprocessing.

Only sleep EEG data prior to awakening (sleep stage S1, S2, S3, S4, and REM) entered the RSA. Following a previously published procedure to detect spontaneous memory processing in sleep ^6^, we averaged power spectra across electrodes within a radius of approximately 3 cm around 32 evenly spread locations of the extended 10–20 system to reduce the dimensionality of the data and to increase the signal-to-noise ratio. To remove amplitude differences between channels, spectra of all channels were then separately normalized between zero and one, which also removes between-subject variability unrelated to the experimental intervention. At the final stage, we applied a spectral sharpening filter to remove the baseline power spectrum by subtracting a moving average of six neighboring frequency bins (window size: 3 Hz) from the signal. This was done to emphasize signal difference ^6^. We then calculated an average PSD of each participant across all sleep segments of the night, separately for REM and NREM sleep, to maximize signal-to-noise ratio. Data between 0.5–30 Hz entered the final analyses ^6^.

### QUANTIFICATION AND STATISTICAL ANALYSIS

#### Representational Similarity Analysis (RSA)

In the present study, we tested whether spontaneous electrical brain activity during sleep holds information about the content of a recently encountered narrative. We employed representational similarity analysis (RSA), a multivariate pattern analysis method that allows comparing how different experimental conditions shape neural activity patterns by assessing the distinctiveness of brain responses across multiple data features ^68^. We used PSD features for similarity calculation because these can integrate information across time and do not depend on specific temporal events. If memory for the story that participants encoded while listening to the audiobook is reactivated during sleep, the narrative should shape brain activity patterns in the sleeping brain, and we should be able to detect information about the narrated content in recordings of electrical brain activity. It can be assumed that brain activity patterns induced by stimulation are similar across participants ^6,25^. We thus hypothesized that brain activity patterns of participants who listened to the same audiobook would be more similar than brain activity pattern of participants who listened to a different audiobook.

##### Leave-one-subject-out RSA

We employed a leave-one-subject-out (LOO) approach for the RSA, correlating brain activity patterns of one participant with the average activity pattern of all other participants who listened to either the same or a different audiobook. This approach gives us a measure of how well an individual’s brain activity patterns conform to the estimated template of brain activity reflecting audiobook reprocessing and can serve as a measure of individual reprocessing strength. We first removed the PSD of one individual (left-out subject) from one audiobook category and averaged the remaining subjects’ PSDs within that audiobook category (**Fig. 1B**). To compute the within-audiobook correlation, we calculated the correlation between the PSD of the left-out subject and the average PSD of all other subjects within that category. The between-audiobook correlations were computed between the PSD of the left-out subject and the averaged PSD of participants in the three other audiobook categories, separately, as shown in **Fig. 1B**. This procedure was repeated for each of the participants. The correlations were implemented using a non-parametric Spearman’s correlation (rho) across the PSD values (1–30 Hz, 0.5 Hz resolution) for all 32 electrode locations (n = 32 x 60), separately for NREM (stages S2, S3, and S4) and REM sleep. We then converted the correlation values to a normal distribution using the inverse hyperbolic tangent (Fisher’s z-transform) and quantified the amount of audiobook-specific neural processing during sleep by contrasting the average within-audiobook correlations with the average between-audiobook correlations. Positive within-between correlation differences (Δcorr) indicate audiobook reprocessing during sleep.

##### Pairwise RSA

In an additional analysis, we re-ran the RSA using the more traditional pairwise correlation approach. Here, we calculated pairwise non-parametric Spearman’s correlations between the average PSDs of all individuals across 32 electrodes, again separately for NREM and REM sleep. We then converted the rho values to a normal distribution using the inverse hyperbolic tangent and averaged all within-audiobook condition correlations, as well as all between audiobook condition correlations that were obtained, to again calculate the within-between difference measure.

##### Permutation testing

To test for statistical significance of the within-between similarity differences, permutation tests were computed separately for all reported analyses. Audiobook labels were shuffled randomly between participants to remove information about audiobook conditions. Then, the exact analysis as reported above was repeated 1000 times, resulting in 1000 average within-between similarity differences that provide the permutation distribution of results when no information about the audiobook is present in the data. The p-value was then computed as the number of averaged within-between differences generated by randomly labeled data that were greater than or equal to the size of the observed average of within-between differences from the correctly labeled data, divided by the number of random permutations and the observed difference (n+1).

##### Searchlight analyses for frequency bands of interest (FOI)

We assessed the contribution of different oscillatory frequencies to memory reprocessing during REM sleep by removing information about the audiobook condition from specific frequency bands and testing its effect on signal similarity differences in the RSA described above. This was done by randomly swapping the features of interest between conditions. We randomly shuffled the data only regarding the FOI, to test whether this significantly decreases informative content. To achieve this, we randomly reassigned the parts of the PSD in REM sleep corresponding to the FOIs between participants, leaving the remaining data structure intact and maintaining correct audiobook labelling for all other frequency bands (delta: 0.5–3.5 Hz, theta: 4–7.5 Hz, alpha: 8–10.5 Hz, beta: 18–30 Hz). We did not include the spindle range (10.5–18 Hz) in our FOI analyses because NREM sleep spindles do not occur during REM sleep. An advantage of this searchlight procedure is that the number of features and the overall data structure are kept constant for all analyses, regardless of the widths of the frequency bands. This procedure was completed 1000 times for each FOI. If the observed within-between similarity difference exceeds 95% of the values in the randomization distribution obtained by shuffling data in the FOI, the respective frequency band can be assumed to hold crucial information about the audiobook and thus significantly contributes to memory processing in sleep. As a control analysis, we also ran a LOO RSA on data within the FOI only, with a reduced number of features that consequently varied between FOIs. This type of analysis regards a limited number of features only, without controlling for the overall structure and dependencies in the data to be analyzed. Results of this supplementary analysis align with the FOI results reported in the main manuscript and are displayed in **Fig. S3.**

##### Searchlight analyses of regions of interest (ROI) in the beta frequency range

After observing audiobook-related processing in REM sleep beta activity, we wanted to investigate the contribution of different brain areas to memory processing in sleep. For this, we performed a region of interest (ROI) analysis. We used whole-brain high density EEG data, analyzing all 128 channels separately, to obtain better regional specificity. In a first step, we confirmed that we can detect audiobook information in sleep if we run the LOO RSA described above on 128 separate channels instead of 32 averaged channels. We replicated our previous findings, showing that the memory is reprocessed in REM sleep, and that REM sleep contains information about the audiobook condition mainly in the beta frequency range (see **Fig. S2**). To investigate regional contributions, we first grouped the EEG channels into six separate regions of interest (ROIs): frontal (n = 34), central (n = 32), temporal (n = 20), parietal (n = 23), and occipital regions (n = 19). All ROIs were bilateral. To test whether specific brain regions had a significant contribution to audiobook processing in the beta range, we permuted the filtered PSD data (18–30 Hz) between participants in these specific ROIs, effectively removing all information regarding the audiobook condition from this part of the data, while keeping the rest of the data correctly labeled. This procedure was completed 1000 times for each ROI and the *p*-values were computed in the same way as described above. Following the same logic, if the within-between difference in correlation values for the completely intact data is significantly higher than the distribution of correlation value differences after permuting information in the beta range for a specific ROI, this region significantly contributes to memory reprocessing during sleep.

#### Behavioral Data Analyses

Statistical analyses were carried out using R (Version 4.3.1., R Core Team (2020)^71^. *Dream reports.* To analyze the content of the dreams, three blind and independent raters received the dream reports in a randomized order. They were asked to indicate which audiobook participants had encoded and to rate how much information about the audiobook they detected in the dream report, as a measure of decision confidence (0 = guessed, 1 = gut feeling, 2 = implicit indication, 3 = explicit indication). The blind raters were highly familiar with the audiobooks. They had listened to the audiobooks and had read the text versions of the books in detail (the text was adapted to match the exact text of the audiobooks). The raters had access both to the actual audiobooks, as well as to the text version of all audiobooks while performing their ratings (with time stamping for each page to allow comparisons between text and audiobook time). They were also provided with the information up to which time point participants had listened to the audiobook and were thus able to compare this to the timestamps in the text version of the audiobook. To compare the rated and actual audiobook, we assessed Cohen’s Kappa in 149 available dream ratings (150 values from 50 reported dreams, one missing rating. Examples of dream reports with a high incorporation rating for each audiobook can be found in the **Table S3**. Note that a chi-square test of independence to examine the relationship between audiobook condition and the correct ratings revealed no significant differences in ratings across audiobook conditions χ²(3, N = 149) = 1.38, *p* = 0.71.

We also performed the same analyses using individual dream reports (n=39), where the audiobook category rated by the blind raters with the highest frequency was selected as a consensus. In cases of disagreement among the three raters, the rating with the highest confidence was chosen. Ambiguous ratings (equal confidence for two different outcomes) were treated as missing values. The similarity between dream reports and audiobooks was compared to all the audiobook passages previously listened to. In the few cases where direct incorporation occurred, information from the segment listened to immediately before that specific awakening was incorporated.

In our concordance analyses, we test whether there is a concordance between rating of all raters and the actual audiobook condition (ground truth), and thus deviates from traditional uses of Cohen’s Kappa in assessing interrater agreement. Specifically, we aim to determine whether the content of dream reports enables blind raters to make correct decisions about what learning material was presented before sleep. The relevant measure is thus not the height of Kappa, but whether the number of corrects rating exceeds the number expected by chance. Thus, in the current study, significance of the concordance analysis indicates that learning material is reactivated in dreams. Please note that raters had to choose an audiobook condition regardless of whether the participants actually had a dream in which incorporated the audiobook (four-way force choice design), such that the base rate of detectable information may be low. Moreover, raters’ decisions were subject to several other noise sources, such as their own memory for the source material. Kappa values measuring agreement between raters and ground truth (actual audiobook condition) are thus expected to be lower than in traditional use cases. The three-way interrater agreements are similarly affected by this and are thus also expected to be low.

##### Audiobook dream reinstatement score

From the ratings on how much information about the audiobook the scorers detected in the dream reports, we computed a separate audiobook dream reinstatement score. The information score was computed by summing up the three raters’ confidence interval by accounting for the blind rater’s correctness individually. More precisely, if a rater was able to correctly identify the audiobook, their information score was multiplied by 1, otherwise by -1. Scores higher than zero thus reflect that information pertaining to the audiobook was incorporated into the dream, allowing correct judgements, with higher scores indicating more direct evidence for incorporation, whereas scores of zero and lower mean that no information about the previously listened audiobook was present in the dream report such that the condition was judged incorrectly. For statistical analyses, we averaged these dream reinstatement scores across all awakenings per individual. We then divided participants into dream incorporators (average value above zero) and dream non-incorporators (average value below zero). A subject-level dream reinstatement score was necessary to relate dream incorporation to participant-specific neural reinstatement scores (see Relating reprocessing strength and dream reinstatement score).

##### Cognitive Measures

Free recall was computed by dividing the number of recalled words by the number of minutes that participants had listened to the audiobook while awake (number of free recall words/minutes audiobook while awake). The word count during free recall ranged from 0 to 470 words (*M* = 88.38, SD = 89.09 words) and the values after adjusting for the time that the participant listened to the audiobook while awake ranged from 0 to 122 (*M* = 11.14, SD = 17.51).

Likewise, audiobook recognition was computed as the percentage of the audiobook passage recognized that was listened to while awake (audiobook time recognized/audiobook runtime while awake). The performance in the audiobook recognition task ranged from 0 to 2675 seconds of audiobook recognized (*M* = 512.2, *SD* = 467.66 seconds), while the percentage values after adjusting for the time ranged from 0 to 1 (*M* = 0.64, SD = 0.30). Since the amount of time that participants spent awake during our repeated awakening paradigm varied depending on how long they took to fall asleep while listening to the audiobook, we also tested whether there were systematic differences in audiobook exposure times across audiobook conditions. We found no significant effect of audiobook condition on time spent awake during the serial awakening paradigm, suggesting that audiobook identity did not systematically bias sleep structure, allowing audiobook classification based on this cofound (F(3, 63) = 0.400, *p* = 0.753).

Note that only audiobook parts played before participants entered sleep were remembered. Participants were not reporting parts that were played during sleep.

For analyses relating memory performance with neural reprocessing and brain activity during sleep, we separately averaged recall and recognition values across sleep awakenings for each participant.

#### Correlation of neural activity with behavioral performance

To relate the strength of neural reprocessing with memory performance, we calculated a neural reprocessing score for each of the participants. This score was quantified by how similar their brain activity during REM sleep was to the activity template gained from the average activity of the other participants in the same audiobook condition (see *Leave-one-subject-out RSA*). We then correlated these values with the individual average recognition and recall performance throughout the night. Similarly, we examined whether EEG activity that reflects audiobook reprocessing during sleep is associated with retention of the content of the audiobook after sleep, we further correlated beta activity in REM sleep with recall and recognition measures. To obtain a measure of beta activity, we averaged the PSD that entered RSA analysis in the beta frequency range (18–30 Hz) for each participant and correlated this measure with the memory performance. We computed the association of beta-activity with a higher word count in dream reports using a Pearson correlation. To delineate the role of beta-activity in memory consolidation, we further conducted a mediation analysis using *mediate* function in R ^70^ to examine whether the number of words recalled from dream reports mediated the relationship between beta activity during REM sleep and the free recall of audiobook content listened to while falling asleep.

#### Relating reprocessing strength and dream reinstatement score

To relate the strength of neural reprocessing with the incorporation of audiobook material into dreams, we calculated a neural reinstatement score for each of the participants. We then compared whether participants who incorporated audiobook information in their dreams showed higher neural reinstatement scores than participants who did not dream of the audiobook.

## References

1. Girardeau, G. (2021). Brain neural patterns and the memory function of sleep. Science 374(*6567*), 560–564. 10.1126/science.abi8370.

2. Klinzing, J.G., Niethard, N., and Born, J. (2019). Mechanisms of systems memory consolidation during sleep. Nat Neurosci. 10.1038/s41593-019-0467-3.

3. Picard-Deland, C., Bernardi, G., Genzel, L., Dresler, M., and Schoch, S.F. (2023). Memory reactivations during sleep: a neural basis of dream experiences? Trends in Cognitive Science.

4. Stickgold, R., Hobson, J.A., Fosse, R., and Fosse, M. (2001). Sleep, learning, and dreams: Off-line memory reprocessing. Science 294, 1052–1057. 10.1126/science.1063530.

5. Nielsen, T.A., and Stenstrom, P. (2005). What are the memory sources of dreaming? Nature 437, 1286–1289. 10.1038/nature04288.

6. Schönauer, M., Alizadeh, S., Jamalabadi, H., Abraham, A., Pawlizki, A., and Gais, S. (2017). Decoding material-specific memory reprocessing during sleep in humans. Nat Commun 8. 10.1038/ncomms15404.

7. Wilson, M.A., and McNaughton, B.L. (1994). Reactivation of hippocampal ensemble memories during sleep. Science 265, 676–679. 10.1126/science.8036517.

8. Peyrache, A., Khamassi, M., Benchenane, K., Wiener, S.I., and Battaglia, F.P. (2009). Replay of rule-learning related neural patterns in the prefrontal cortex during sleep. Nature Neuroscience 2009 12:7 12, 919–926. 10.1038/nn.2337.

9. Schreiner, T., Petzka, M., Staudigl, T., and Staresina, B.P. (2021). Endogenous memory reactivation during sleep in humans is clocked by slow oscillation-spindle complexes. Nat Commun 12. 10.1038/s41467-021-23520-2.

10. Louie, K., and Wilson, M.A. (2001). Temporally Structured Replay of Awake Hippocampal Ensemble Activity during Rapid Eye Movement Sleep. Neuron 29, 145–156. 10.1016/S0896-6273(01)00186-6.

11. Klepel, F., and Schredl, M. (2019). Correlation of task-related dream content with memory performance of a film task – A pilot study. International Journal of Dream Research 12, 112–118. 10.11588/ijodr.2019.1.59320.

12. Siegel, J.M. (2001). The REM sleep-memory consolidation hypothesis. Science 294, 1058–1063. 10.1126/SCIENCE.1063049/ASSET/164F2988-F16E-42CE-AB90-2B196329FAD8/ASSETS/GRAPHIC/SE4319896001.JPEG.

13. Ji, D., and Wilson, M.A. (2007). Coordinated memory replay in the visual cortex and hippocampus during sleep. Nat Neurosci 10, 100–107. 10.1038/nn1825.

14. Kumar, D., Koyanagi, I., Carrier-Ruiz, A., Vergara, P., Srinivasan, S., Sugaya, Y., Kasuya, M., Yu, T.S., Vogt, K.E., Muratani, M., et al. (2020). Sparse Activity of Hippocampal Adult-Born Neurons during REM Sleep Is Necessary for Memory Consolidation. Neuron 107, 552–565.e10. 10.1016/j.neuron.2020.05.008.

15. Maquet, P., Laureys, S., Peigneux, P., Fuchs, S., Petiau, C., Phillips, C., Aerts, J., Del Fiore, G., Degueldre, C., Meulemans, T., et al. (2000). Experience-dependent changes in changes in cerebral activation during human REM sleep. Nat Neurosci 3, 831–836. 10.1038/77744.

16. Zhang, H., Fell, J., and Axmacher, N. (2018). Electrophysiological mechanisms of human memory consolidation. Nat Commun 9. 10.1038/s41467-018-06553-y.

17. Rasch, B., Büchel, C., Gais, S., and Born, J. (2007). Odor cues during slow-wave sleep prompt declarative memory consolidation. Science 315, 1426–1429. 10.1126/science.1138581.

18. Antony, J.W., Gobel, E.W., O’Hare, J.K., Reber, P.J., and Paller, K.A. (2012). Cued memory reactivation during sleep influences skill learning. Nat Neurosci 15, 1114–1116. 10.1038/nn.3152.

19. Abdellahi, M.E., Koopman, A.C., Treder, M.S., and Lewis, P.A. (2023). Targeted memory reactivation in human REM sleep elicits detectable reactivation. Elife.

20. Stickgold, R., Malia, A., Maguire, D., Roddenberry, D., and O’Connor, M. (2000). Replaying the game: Hypnagogic images in normals and amnesics. Science 290, 350–353. 10.1126/science.290.5490.350.

21. Cipolli, C., Fagioli, I., Mazzetti, M., and Tuozzi, G. (2004). Incorporation of presleep stimuli into dream contents: Evidence for a consolidation effect on declarative knowledge during REM sleep? J Sleep Res 13, 317–326. 10.1111/J.1365-2869.2004.00420.X.

22. Wamsley, E.J., Tucker, M., Payne, J.D., Benavides, J.A., and Stickgold, R. (2010). Dreaming of a Learning Task Is Associated with Enhanced Sleep-Dependent Memory Consolidation. Current Biology 20, 850–855. 10.1016/j.cub.2010.03.027.

23. Wamsley, E.J., and Stickgold, | Robert (2018). Dreaming of a learning task is associated with enhanced memory consolidation: Replication in an overnight sleep study. Wiley Online Library 28. 10.1111/jsr.12749.

24. Schoch, S.F., Cordi, M.J., Schredl, M., and Rasch, B. (2019). The effect of dream report collection and dream incorporation on memory consolidation during sleep. J Sleep Res 28. 10.1111/jsr.12754.

25. Chen, J., Leong, Y.C., Honey, C.J., Yong, C.H., Norman, K.A., and Hasson, U. (2017). Shared memories reveal shared structure in neural activity across individuals. Nat Neurosci 20, 115–125. 10.1038/nn.4450.

26. Baldassano, C., Hasson, U., and Norman, K.A. (2018). Representation of real-world event schemas during narrative perception. Journal of Neuroscience 38, 9689–9699. 10.1523/JNEUROSCI.0251-18.2018.

27. Cairney, S.A., Guttesen, A.á. V., El Marj, N., and Staresina, B.P. (2018). Memory Consolidation Is Linked to Spindle-Mediated Information Processing during Sleep. Current Biology 28, 948–954.e4. 10.1016/j.cub.2018.01.087.

28. Salvesen, L., Capriglia, E., Dresler, M., and Bernardi, G. (2023). Influencing dreams through sensory stimulation: a systematic review 10.1016/j.tics.2023.02.006.

29. Siclari, F., Baird, B., Perogamvros, L., Bernardi, G., LaRocque, J.J., Riedner, B., Boly, M., Postle, B.R., and Tononi, G. (2017). The neural correlates of dreaming. Nat Neurosci 20, 872–878. 10.1038/nn.4545.

30. Bellana, B., Mahabal, A., and Honey, C.J. (2022). Narrative thinking lingers in spontaneous thought. Nature Communications 2022 13:1 *13*, 1–16. 10.1038/s41467-022-32113-6.

31. Schredl, M., and Hofmann, F. (2003). Continuity between waking activities and dream activities. Conscious Cogn 12, 298–308. 10.1016/S1053-8100(02)00072-7.

32. Empson, J.A., and Clarke, P.R. (1970). Rapid Eye Movements and Remembering. Nature. 10.1038/227287a0.

33. Tilley, A.J., and Empson, J.A.C. (1978). REM sleep and memory consolidation. Biol Psychol 6, 293–300. 10.1016/0301-0511(78)90031-5.

34. Smith, C.T., Nixon, M.R., and Nader, R.S. (2004). Posttraining increases in REM sleep intensity implicate REM sleep in memory processing and provide a biological marker of learning potential. Learning and Memory 11, 714–719. 10.1101/lm.74904.

35. Poe, G.R., Nitz, D.A., McNaughton, B.L., and Barnes, C.A. (2000). Experience-dependent phase-reversal of hippocampal neuron firing during REM sleep. Brain Res 855, 176–180. 10.1016/S0006-8993(99)02310-0.

36. Mandai, O., Guerrien, A., Sockeel, P., Dujardin, K., and Leconte, P. (1989). REM sleep modifications following a Morse code learning session in humans. Physiol Behav 46, 639–642. 10.1016/0031-9384(89)90344-2.

37. Fischer, S., Hallschmid, M., Elsner, A.L., and Born, J. (2002). Sleep forms memory for finger skills. Proc Natl Acad Sci U S A 99, 11987–11991. 10.1073/PNAS.182178199.

38. McDevitt, E.A., Duggan, K.A., and Mednick, S.C. (2015). REM sleep rescues learning from interference. Neurobiol Learn Mem 122, 51–62. 10.1016/J.NLM.2014.11.015.

39. McDevitt, E.A., Kim, G., Turk-Browne, N.B., and Norman, K.A. (2024). The role of REM sleep in neural differentiation of memories in the hippocampus. Preprint, 10.1101/2024.11.01.621588 https://doi.org/10.1101/2024.11.01.621588.

40. Peigneux, P., Laureys, S., Fuchs, S., Destrebecqz, A., Collette, F., Delbeuck, X., Phillips, C., Aerts, J., Del Fiore, G., Degueldre, C., et al. (2003). Learned material content and acquisition level modulate cerebral reactivation during posttraining rapid-eye-movements sleep. Neuroimage 20, 125–134. 10.1016/S1053-8119(03)00278-7.

41. Choi1 Jisoo, Carmichael, J., Williams, S., and Etter, G. (2024). Structured and unstructured reactivations during REM sleep are modulated by novel experiences. bioRxiv. 10.1101/2024.09.05.611474.

42. Bollmann, L., Baracskay, P., Stella, F., and Csicsvari, J. (2023). Sleep stages antagonistically modulate reactivation drift. bioRxiv, 2023.10.13.562165. 10.1101/2023.10.13.562165.

43. Smith, C. (2001). Sleep states and memory processes in humans: Procedural versus declarative memory systems. Sleep Med Rev 5, 491–506. 10.1053/smrv.2001.0164.

44. Izawa, S., Chowdhury, S., Miyazaki, T., Mukai, Y., Ono, D., Inoue, R., Ohmura, Y., Mizoguchi, H., Kimura, K., Yoshioka, M., et al. (2019). REM sleep-active MCH neurons are involved in forgetting hippocampus-dependent memories.

45. Poe, G.R. (2017). Sleep is for forgetting. Journal of Neuroscience 37, 464–473. 10.1523/JNEUROSCI.0820-16.2017.

46. Schreiner, T., Doeller, C.F., Jensen, O., Rasch, B., and Staudigl, T. (2018). Theta Phase-Coordinated Memory Reactivation Reoccurs in a Slow-Oscillatory Rhythm during NREM Sleep. Cell Rep 25, 296–301. 10.1016/j.celrep.2018.09.037.

47. Wang, B., Antony, J.W., Lurie, S., Brooks, P.P., Paller, K.A., and Norman, K.A. (2019). Targeted Memory Reactivation during Sleep Elicits Neural Signals Related to Learning Content. J Neurosci 39, 6728–6736. 10.1523/JNEUROSCI.2798-18.2019.

48. Cairney, S.A., Guttesen, A.á. V., El Marj, N., and Staresina, B.P. (2018). Memory Consolidation Is Linked to Spindle-Mediated Information Processing during Sleep. Current Biology 28, 948–954.e4. 10.1016/j.cub.2018.01.087.

49. Liu, J., Xia, T., Chen, D., Yao, Z., Zhu, M., Antony, J.W., Lee, T.M.C., and Hu, X. (2023). Item-specific neural representations during human sleep support long-term memory. PLoS Biol 21, e3002399. 10.1371/JOURNAL.PBIO.3002399.

50. Williamson, P. c., Csima, A., Galin, H., and Mamelak, M. (1986). Spectral EEG correlates of dream recall. Biol Psychiatry 21, 717–723. 10.1016/0006-3223(86)90236-2.

51. Chellappa, S.L., Frey, S., Knoblauch, V., and Cajochen, C. (2011). Cortical activation patterns herald successful dream recall after NREM and REM sleep. Biol Psychol 87, 251–256. 10.1016/j.biopsycho.2011.03.004.

52. Morton, N.W., and Polyn, S.M. (2017). Beta-band activity represents the recent past during episodic encoding. Neuroimage 147, 692–702. 10.1016/J.NEUROIMAGE.2016.12.049.

53. Jayachandran, M., Viena, T.D., Garcia, A., Veliz, A.V., Leyva, S., Roldan, V., Vertes, R.P., and Allen, T.A. (2023). Nucleus reuniens transiently synchronizes memory networks at beta frequencies. Nat Commun 14, 4326. 10.1038/s41467-023-40044-z.

54. Ferri, R., Cosentino, F.I.I., Elia, M., Musumeci, S.A., Marinig, R., and Bergonzi, P. (2001). Relationship between Delta, Sigma, Beta, and Gamma EEG bands at REM sleep onset and REM sleep end. Clinical Neurophysiology 112, 2046–2052. 10.1016/S1388-2457(01)00656-3.

55. Pesonen, A.K., Makkonen, T., Elovainio, M., Halonen, R., Räikkönen, K., and Kuula, L. (2021). Presleep physiological stress is associated with a higher cortical arousal in sleep and more consolidated REM sleep. Stress 24, 667–675. 10.1080/10253890.2020.1869936;WGROUP:STRING:PUBLICATION.

56. Freedman, R.R. (1986). EEG power spectra in sleep-onset insomnia. Electroencephalogr Clin Neurophysiol 63, 408–413. 10.1016/0013-4694(86)90122-7.

57. Kuo, T.B.J., Chen, C.Y., Hsu, Y.C., and Yang, C.C.H. (2016). EEG beta power and heart rate variability describe the association between cortical and autonomic arousals across sleep. Autonomic Neuroscience 194, 32–37. 10.1016/J.AUTNEU.2015.12.001.

58. Speth, C., and Speth, J. (2016). The borderlands of waking: Quantifying the transition from reflective thought to hallucination in sleep onset. Conscious Cogn 41, 57–63. 10.1016/j.concog.2016.01.009.

59. Horikawa, T., Tamaki, M., Miyawaki, Y., and Kamitani, Y. (2013). Neural decoding of visual imagery during sleep. Science 340, 639–642. 10.1126/science.1234330.

60. Siclari, F., LaRocque, J.J., Postle, B.R., and Tononi, G. (2013). Assessing sleep consciousness within subjects using a serial awakening paradigm. Front Psychol 4. 10.3389/fpsyg.2013.00542.

61. Konkoly, K.R., Appel, K., Chabani, E., Mangiaruga, A., Gott, J., Mallett, R., Caughran, B., Witkowski, S., Whitmore, N.W., Mazurek, C.Y., et al. (2021). Real-time dialogue between experimenters and dreamers during REM sleep. Current Biology 31, 1417–1427.e6. 10.1016/J.CUB.2021.01.026/ATTACHMENT/03D032D8-0FCC-476C-8BD5-23220137719C/MMC2.PDF.

62. Wong, W., Cristine Andrade, K., Andrillon, T., Barros de Araujo, D., Arnulf, I., Avvenuti, G., Baird, B., Bellesi, M., Bergamo, D., Bernardi, G., et al. DREAM: A Dream EEG and Mentation database 1 DREAM: A Dream EEG and 1 Mentation database 2.

63. Wynn, J.S., Shen, K., and Ryan, J.D. (2019). Eye Movements Actively Reinstate Spatiotemporal Mnemonic Content. Vision 2019, Vol. 3, Page 21 3, 21. 10.3390/VISION3020021.

64. Andrillon, T., Nir, Y., Cirelli, C., Tononi, G., and Fried, I. (2015). Single-neuron activity and eye movements during human REM sleep and awake vision. Nature Communications 2015 6:1 6, 1–10. 10.1038/ncomms8884.

65. Maranci, J.B., Nigam, M., Masset, L., Msika, E.F., Vionnet, M.C., Chaumereil, C., Vidailhet, M., Leu-Semenescu, S., and Arnulf, I. (2022). Eye movement patterns correlate with overt emotional behaviours in rapid eye movement sleep. Scientific Reports 2022 12:1 12, 1–9. 10.1038/s41598-022-05905-5.

66. Senzai, Y., and Scanziani, M. (2022). A cognitive process occurring during sleep is revealed by rapid eye movements. Science 377, 999–1004. 10.1126/science.abp8852.

67. Delorme, A., and Makeig, S. (2004). EEGLAB: an open sorce toolbox for analysis of single-trail EEG dynamics including independent component anlaysis. J Neurosci Methods 134, 9–21. 10.1016/j.jneumeth.2003.10.009.

68. Diedrichsen, J., Kriegeskorte, N., Rn Diedrichsen, J., and Kriegeskorte, N. (2017). Representational models: A common framework for understanding encoding, pattern-component, and representational-similarity analysis. PLoS Comput Biol 13. 10.1371/journal.pcbi.1005508.

69. Siclari, F., Bernardi, G., Cataldi, J., and Tononi, G. (2018). Dreaming in NREM sleep: A high-density EEG study of slow waves and spindles. Journal of Neuroscience 38, 9175–9185. 10.1523/JNEUROSCI.0855-18.2018.

70. Imai, K., Keele, L., Tingley, D., and Yamamoto, T. (2010). Causal Mediation Analysis Using R. 129–154. 10.1007/978-1-4419-1764-5_8.

71. Team, R.C. (2013). R: A language and environment for statistical computing.

