## Supplemental Figures for "Pre-sleep experiences shape neural activity and dream content in the sleeping brain"

### Supplemental Information

**Fig. S1. Neural reinstatement using pairwise instead of leave-one-out (LOO) representational similarity analyses (related to Fig. 2). A**. We calculated a non-parametric Spearman's correlation between power spectral density values (PSD, dimensions: 32 channel x 60 Hz) of individuals (k=67 awakening from 19 participants). Note that the last 250 trials of each sleep period before awakening entered these analyses, corresponding to the last quarter hour of sleep (16.7 minutes). We then converted the obtained rho values to a normal distribution using Fisher's inverse hyperbolic tangent transform**. B.** Representational similarity analyses (RSA) results for REM and NREM sleep. We converted correlation values (rho) to a normal distribution using the Fisher inverse hyperbolic tangent transformation. The histograms represent the permutation distribution of within-between differences for randomly relabeled data (x-axis; Δcorr). Representational similarity matrices are displayed to the right. Upper triangle: entries on the diagonal of the similarity matrix denoted by black lines represent pairwise within-audiobook correlations, off-diagonal entries show pairwise between-audiobook correlations. Lower triangle: average within (diagonal) and between (off-diagonal) correlation values for each audiobook condition. REM: within-audiobook correlation $M = 0.312$, between-audiobook correlation $M = 0.275$; NREM: within-audiobook correlation $M = 0.393$, between-audiobook correlation $M = 0.409$. Note that a total of 67 awakenings from 19 participants entered the RSA.

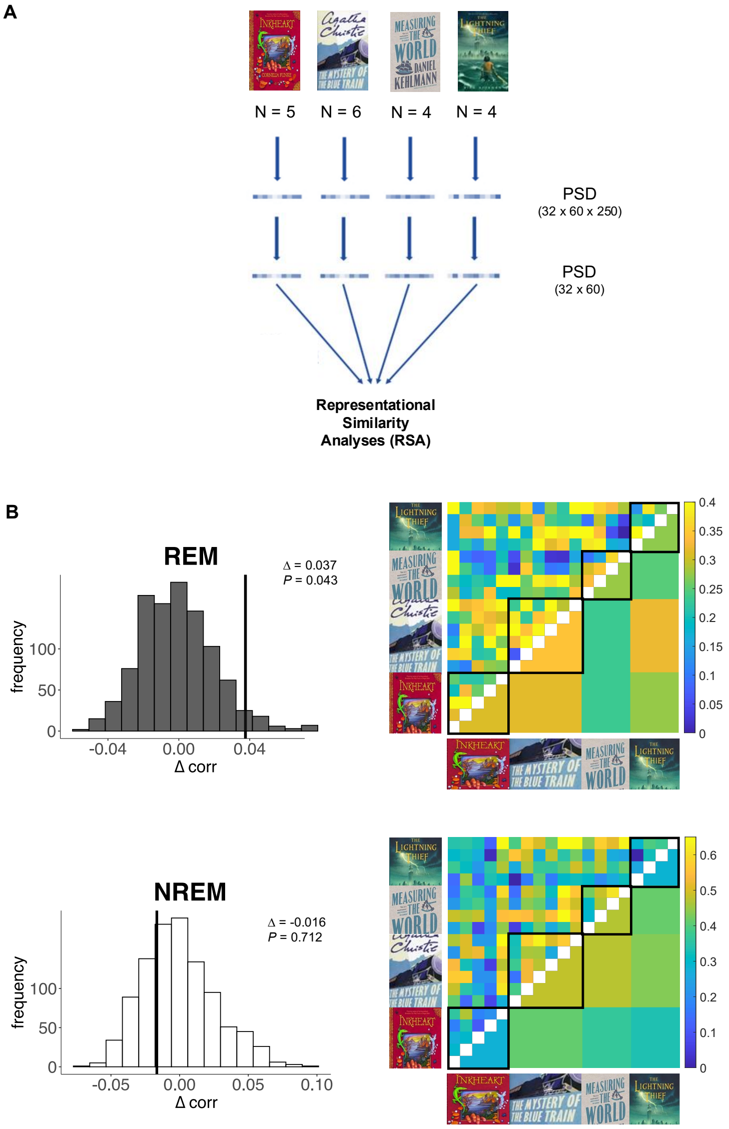

**Fig. S2. Neural reinstatement using whole brain EEG with all 128 individual channels instead of 32 averaged channels (related to Fig. 2).** The histograms represent the permutation distribution of within-between differences for randomly relabeled data (x-axis; Δcorr) computed using the leave-one-subject-out representational similarity analysis (RSA) approach, separately for **A.** NREM (Δcorr = -0.040, *P* = 0.747) and **B.** REM sleep (Δcorr = -0.040, *P* = 0.047). **C**. To assess the contribution of the different oscillatory activity to memory processing in REM sleep, we shuffled power spectral density (PSD) patterns between participants in specific frequency bands, thus removing audiobook information from these parts of the data while keeping the correct audiobook labels intact for all other frequency bands. The vertical black lines indicate the observed Δcorr for REM sleep with all frequency ranges intact. REM sleep beta activity (18–30 Hz) contributes significantly to the strength of neural reprocessing. Note that a total of 67 awakenings from 19 participants entered the RSA.

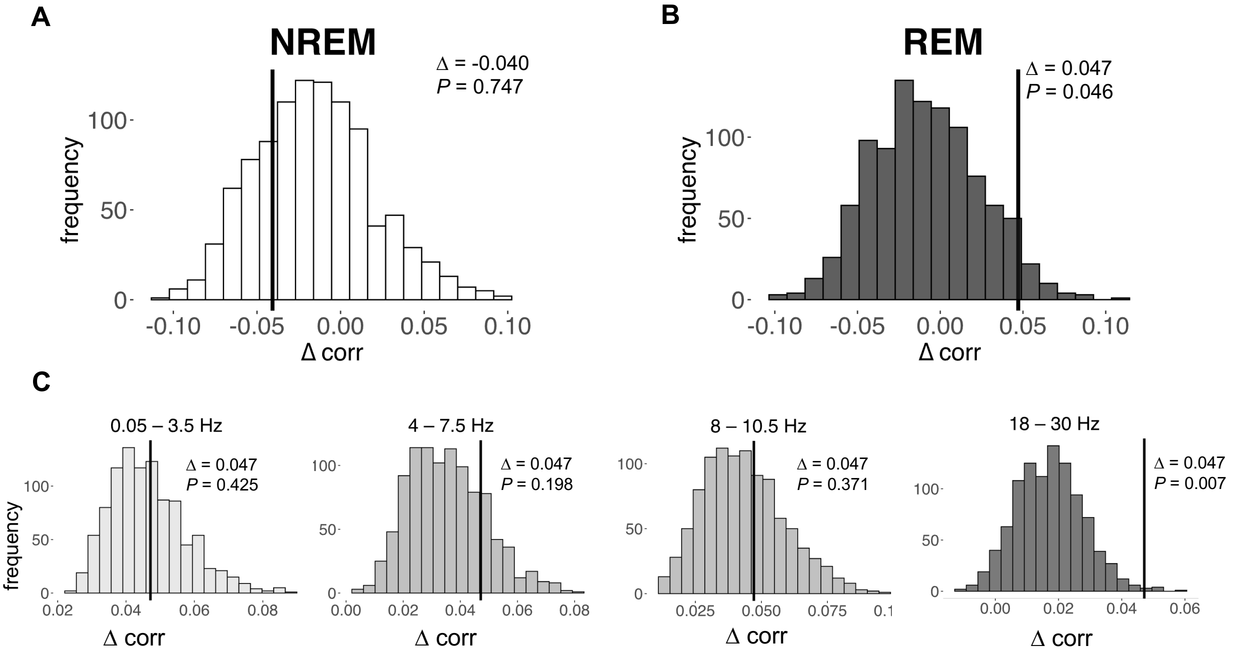

**Fig. S3. The contribution of individual frequency bands to neural reinstatement (related to Fig. 3).** Only data in individual frequency bands was included in this analysis. The histograms represent the permutation distribution of random within-between differences (x-axis; Δcorr), while the black line indicates the observed differences. To assess the contribution of the different frequencies to RSA during REM sleep, we shuffled individuals' frequency bands (delta: 0.5–3.5 Hz, theta: 4–7.5 Hz, alpha: 8–10.5 Hz, beta: 18–30 Hz, *frequency of interest* (FOI); shown in dot line). Only beta frequency (18–30 Hz) contributes significantly to the neural reinstatement. Note that a total of 67 awakenings from 19 participants entered the representational similarity analyses.

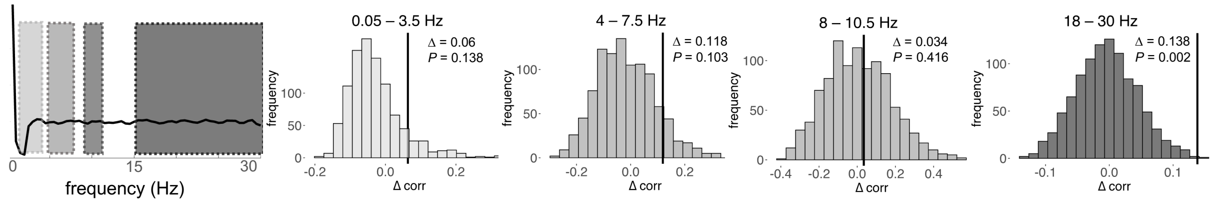

**Fig. S4. Classified dream content separately for REM and NREM sleep (related to Fig. 5).** Cohen's Kappa was used to compare the rated and actual audiobooks, separately for REM and NREM sleep. We observe that the blind raters were able to judge which audiobook participants encoded based on the content of subsequent dreams when the participants were awoken in **A.** REM sleep (κ = 0.343, z = 2.88, *P* = 0.003, percent correct ratings = 54.2%), but not in **B.** NREM sleep (stages S2, S3, S4; κ = 0.08, z = 1.43, *P* = 0.154, percent correct ratings = 31.5%). Note that some participants were awoken from S1 sleep (k=15) or were already awake (k=18) when we entered the sleep chambers to record dream reports such that the combined number of REM sleep and NREM sleep dream ratings is lower than the available total of k=149 dream ratings reported in **Fig. 5.**

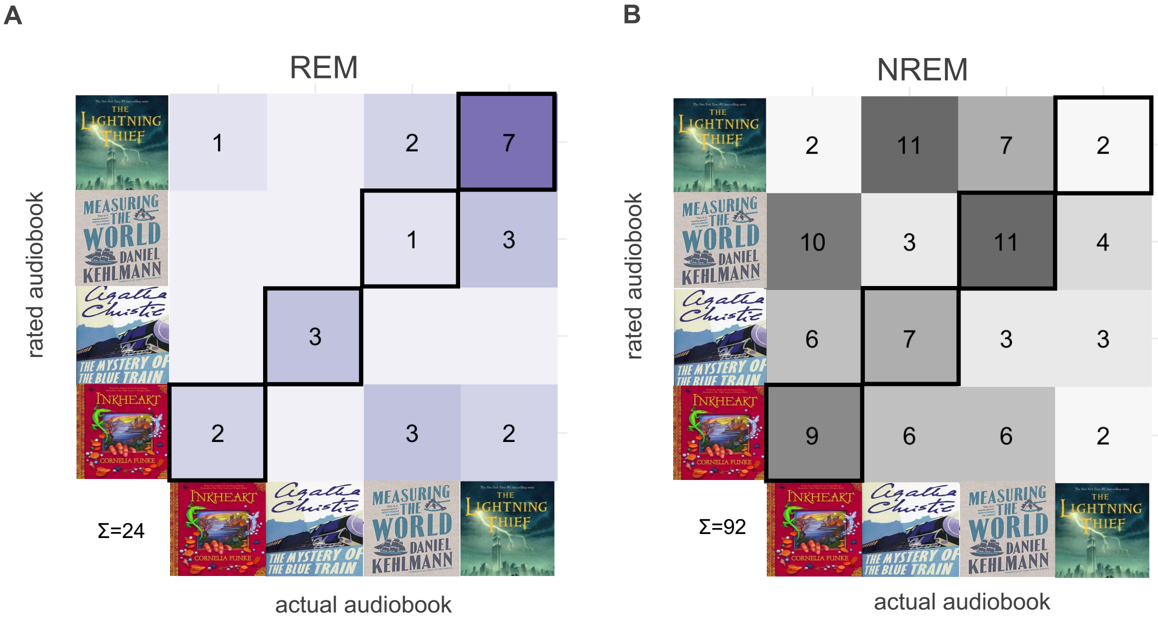

**Fig. S5. Data per sleep stage and participant used in representational similarity analysis (related to EEG Data Preprocesing).** The x-axis represents the total duration (in minutes) across all awakenings, while the y-axis represents the sleep stages from S1 to REM. Each graph corresponds to an individual participant, who listened to one of the following audiobooks: Inkheart (Participants #2, 3, 4, 10, 15, shown in red), The Mystery of the Blue Train (Participants #5, 11, 12, 14, 16, 19, shown in blue), Measuring the World (Participants #1, 6, 8, 13, shown in gray), or Percy Jackson & the Olympians (Participants #7, 9, 17, 18, shown in blue).
Note that NREM sleep consists of concatenated stages S2, S3, and S4.

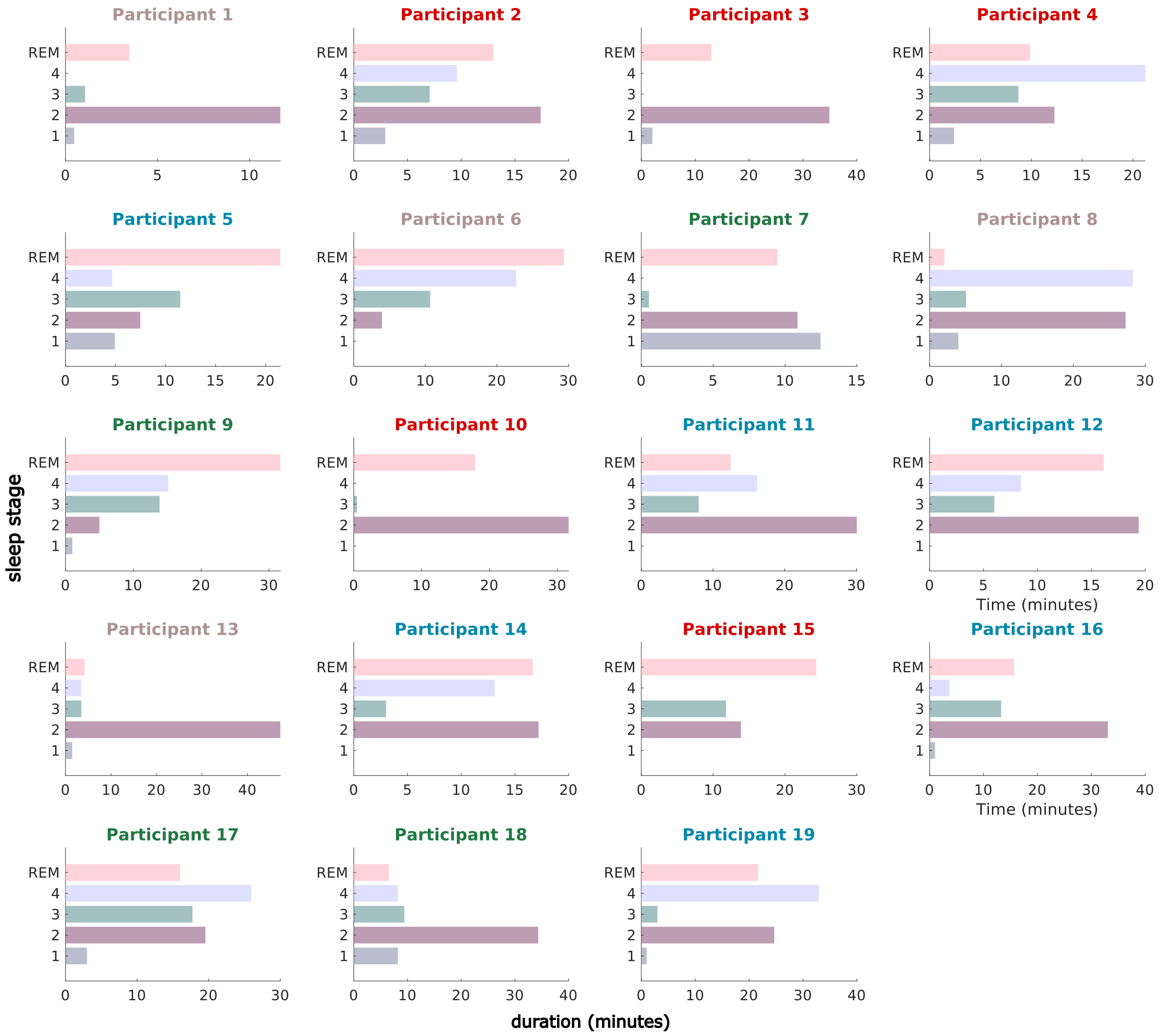

**Table S1. Averaged sleep data in minutes (mean ± SD; related to EEG Recording).**

| | time to fall a sleep | S1 | S2 | S3 | S4 | REM | TST |
|---|---|---|---|---|---|---|---|
| SP 1 | 21 ± 18.8 | 7.1 ± 4.6 | 19.6 ± 13.7 | 11.7 ± 6.7 | 25.9 ± 24 | 9.7 ± 0.8 | 73.9 ± 7.8 |
| SP 2 | 20.1 ± 18.2 | 8.2 ± 4.5 | 34.8 ± 15.6 | 10.7 ± 7.7 | 9.6 ± 9.8 | 16.8 ± 18.1 | 80 ± 11 |
| SP 3 | 21.3 ± 13.8 | 18 ± 33.5 | 46.7 ± 20.7 | 8.3 ± 5.7 | 9.4 ± 5 | 23.6 ± 14.7 | 106 ± 15.6 |
| SP 4 | 14.8 ± 10.8 | 9.9 ± 6.3 | 39.7 ± 22 | 10.6 ± 10.2 | 7.8 ± 7.6 | 31.4 ± 17.1 | 99.5 ± 14.7 |
| SP 5 | 9 ± 9.5 | 5.3 ± 3.3 | 12.7 ± 10.8 | 8.9* | 20.5* | 38 ± 26.9 | 85.5 ± 13 |

*SP = Sleep Period, TST = Total Sleep Time,*

*Note.* Only one participant reached S3/S4 in the 5th sleep segment.

**Table S2. Descriptive statistics of NREM and REM trials before each awakening (Related to EEG Data Preprocessing)**.

| Stage | awakening | count | mean | sd | median | min | max |
|---|---|---|---|---|---|---|---|
| NREM | 1 | 16 | 14.97 | 3.38 | 16.67 | 7.20 | 16.67 |
| NREM | 2 | 16 | 12.25 | 4.40 | 12.67 | 3.67 | 16.67 |
| NREM | 3 | 18 | 10.67 | 6.52 | 12.97 | 0.00 | 16.67 |
| NREM | 4 | 14 | 9.03 | 6.94 | 6.63 | 0.00 | 16.67 |
| NREM | 5 | 3 | 5.56 | 9.62 | 0.00 | 0.00 | 16.67 |
| REM | 1 | 16 | 1.12 | 3.08 | 0.00 | 0.00 | 9.47 |
| REM | 2 | 16 | 3.85 | 4.75 | 2.00 | 0.00 | 13.00 |
| REM | 3 | 18 | 4.58 | 6.46 | 0.23 | 0.00 | 16.13 |
| REM | 4 | 14 | 6.49 | 6.82 | 5.10 | 0.00 | 16.67 |
| REM | 5 | 3 | 11.11 | 9.62 | 16.67 | 0.00 | 16.67 |

*Note.* The table reports the duration (in min) per sleep stage (NREM, REM) used for representation similarity analysis. While trial counts may vary across participants and awakenings, durations were averaged across awakenings for analysis.

**Table S3. Dream reports (Related to Fig 4, Behavioral Data Analyses).**

| | | | |
|---|---|---|---|
| 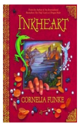 | "I was in an airplane with my father and he was still driving on the runway, then he said: "I have to take off now" or something. And then he just drove faster but somehow he couldn't take off. He let the plane slow to a stop. I had asked him: "But, you can't actually fly" and then he said "Let's see". it was at night and it was raining and there were people on the runway and they jumped away from him." | 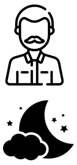 | *father, night* |
| 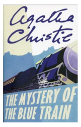 | "What went through my mind was something about stolen diamonds" | 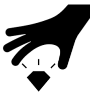 | *theft of jewels* |
| 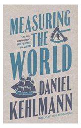 | "Someone said to me that an exchange to Brazil doesn't make sense, I should only go to Portugal; I dreamt that my experimenter specifically showed it to me on the iPhone on a map" | 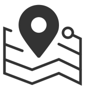 | *travel locations similar to Humboldt's (South America)* |
| 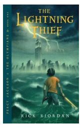 | "I had a huge fight with my boyfriend; we were running around the flat and yelling at each other; there was water damage in the flat." | 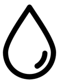 | *water, fountain were incorporated into apartment* |

***Note.*** Exemplary dream reports for which raters scored either direct or indirect incorporations of audiobook content. Comments by the raters are printed on the right. Many sections of Inkheart are set at night – the protagonist's father is a prominent character in the book who can read people into stories (magical skills). Diamonds are stolen in The Mystery of the Blue Train. In Measuring the World, Humboldt imbarks on travels with destinations (South America) similar to those that surfaced in the dream below, maps are used as a tool for navigating. In Percy Jackson, The Lightning Thief, water features as a main element of magic, Percy is the child of Poseidon (the God of the Seas), and a fountain features in the narrative.
